# Arboviruses disrupt salivary gland organization and decrease salivation in *Aedes aegypti* mosquitoes

**DOI:** 10.1101/2025.05.16.654423

**Authors:** Lucie Souriac, Mateus Berni, Benjamin Voisin, Carlos F. Estevez-Castro, Marine Le Berder, Roenick P. Olmo, Thiago H.J.F. Leite, Hacène Medkour, Julien Pompon, Eric Marois, Fabien Aubry, João T. Marques

## Abstract

*Aedes aegypti* mosquitoes are major vectors for arboviruses. Virus transmission occurs during blood feeding, when infected mosquitoes release both viral particles and saliva. Although saliva proteins can influence viral transmission, their study has been limited by the difficulty of quantifying salivation in natural transmission contexts. Here, we generated a transgenic *Aedes aegypti* line that secretes fluorescent markers into saliva, enabling direct visualization and quantification of salivation. Using this tool, we found that secretion of saliva reporters is robust across diverse physiological conditions but is markedly reduced following Zika or chikungunya virus infection, both in artificial salivation and mouse feeding assays. In the salivary glands, viral antigens displaced saliva reporters in secretory acinar cells, suggesting a segregation mechanism by which the virus is released independently from salivation. Importantly, diminished release of saliva proteins may contribute to the increased probing behavior observed in infected mosquitoes, which can enhance the chances of virus transmission. Together, these findings suggest that virus-induced suppression of salivation may be an adaptive feature that enhances transmission efficiency.

## Introduction

Every year worldwide, mosquitoes are responsible for millions of human deaths due to the pathogens they vector. Arboviruses such as yellow fever (YFV), dengue (DENV), Zika (ZIKV) or chikungunya (CHIKV) are transmitted by *Aedes* mosquitoes. Current global warming and urbanization dynamics facilitate the spread of mosquitoes in temperate regions, increasing the risk of arboviruses circulation in these areas^1,2^. If no efficient measures are taken, 60% of the world population is expected to be at risk to contract DENV by the year 2080^2^. The main vector of arboviruses, *Ae. aegypti*, is well adapted to urban landscapes and feeds preferentially on humans^3,4^. It is established in tropical and subtropical regions of the world, allowing the endemic spread of mosquito-borne viruses in these regions^5–8^.

Mosquitoes get infected when they blood feed on a viremic vertebrate host. Viruses acquired in the blood infect and replicate in enterocytes in the midgut before disseminating to the hemocoel of the mosquito^9^. Once in the circulation of the mosquito, the virus will infect and replicate in the salivary glands^10^. There, the virus accumulates in specific sites at the apical cavity of acinar cells that appear distinct from vesicles containing saliva^11^. There, the virus is released when the mosquito probes the skin of the next host to blood feed again. Since the virus is released from the salivary glands, it is assumed that transmission is intimately associated with the release of saliva proteins by the mosquito^12^. Saliva has an important role in modulating different aspects of blood feeding on a vertebrate host, being involved in the inhibition of host responses as well as feedback loops that fine-tune mosquito behavior^12^. For example, LIPS proteins in the saliva modify proboscis morphology and proprioception signaling to promote feeding on a vertebrate host^13^. Of note, saliva proteins have been shown to affect arbovirus infections. For example, venom allergen-1 and sialokinin increase the pathogenicity of ZIKV by activating autophagy or vascular leakage at the biting site, respectively^14,15^. In contrast, the D7 saliva protein interacts directly with DENV particles, inhibiting infection^16^, and aegyptin activates immune responses in mice, decreasing DENV infection^17^. Natural transmission by the mosquito vector affects the pathogenesis of arbovirus infections compared to artificial injection of the virus^18,19^. These results support a role for mosquito saliva in modulating arbovirus infection in the mammalian host. However, natural infection models with infected mosquitoes have not directly measured the release of saliva^18,19^. Other studies based on co-injection of saliva or salivary gland extracts with the virus assume this is what happens during natural transmission by infected mosquitoes^20,21^. Indeed, tracking the release of saliva proteins and virus by an infected mosquito is a challenge. Some studies have assayed viral transmission using forced salivation, where mosquitoes are immobilized and allowed to salivate in drops of oil^22,23^ or tips filled with physiological buffers^24^. These studies provided indirect estimations of the release of saliva proteins analyzing the volume of saliva released or engorgement of the mosquito. In forced salivation assays, a disconnect between the volume of saliva and virus released from mosquitoes has been reported^25^. It also remains unclear whether there is a direct correlation between volume of liquid and the amount of saliva proteins released. These may also help explain why forced salivation assays often underestimate the amount of transmitted virus compared to natural feeding on mice^26,27^. Overall, technical limitations have hindered understanding the connection between salivation (e.g. release of saliva proteins) and virus transmission.

To address these important questions in the field, we developed a transgenic *Ae. aegypti* reporter line expressing epitope-tagged fluorescent proteins in the distal-lateral lobes of the salivary glands. We also introduced signal peptides to these fluorescent proteins that direct their secretion in the saliva. Using this transgenic line, we were able to quantify the release of saliva reporters at the resolution of individual mosquitoes and to characterize basic aspects of the physiology of salivation. Finally, it also allowed us to study the correlation between salivation and transmission of two relevant arboviruses, ZIKV and CHIKV.

## Results

### Generating a transgenic mosquito line to track salivation

To generate a mosquito line that expresses a reporter for salivation, we first searched for genes encoding proteins secreted within the saliva. The *30Ka* (*AAEL010228* encoding SAAG-4) and *30Kb* (*AAEL010235* encoding aegyptin) genes are controlled by a single bidirectional promoter that drives their expression in the distal-lateral lobes of *Ae. aegypti* salivary glands^28,29^. The *30Kb* gene is also expressed in the proximal-lateral lobes to a lesser extent^28^. Using the recent Mosquito Cell Atlas^30^, we confirmed that *30Ka* and *30Kb* are specifically expressed in the distal-lateral lobes of the salivary gland of *Ae. aegypti* mosquitoes and the proximal lobes in the case of *30Kb* (**Figure 1a,b**). They were among the top 35 expressed transcripts in the distal-lateral lobes of the salivary glands (**Supplementary table 1**). Significant expression of *30Ka* and *30Kb* was not detected in other mosquito tissues (**Supplementary figure 1a**).The SAAG-4 and aegyptin proteins, encoded by *30Ka* and *30Kb* genes, respectively, are among the top 20 most abundant secreted in the saliva^31,32^. Mass spectrometry analyses show that SAAG-4 and aegyptin are also in the top 10 proteins found in salivary gland extracts^16^. Of note, the *30K* promoter has been used to successfully drive specific transgene expression in the salivary gland of mosquitoes^33,34^, but not to track salivation. Here we built a construct with the *30K* promoter maintaining the signal peptides encoded by *30Ka* and *30Kb* genes in fusion with the fluorescent markers mTurquoise2 (mTurq) and enhanced Yellow Fluorescent Protein (YFP), respectively (**Figure 1c**). V5 and HA epitope tags were also inserted to facilitate detection of mTurq and YFP, respectively (**Figure 1c**). DsRed2 was used as a marker for transgenesis driven by a ubiquitous promoter (*AAEL003888*, *polyubiquitin*, *PUb*) on the same *piggyBac* construct (**Figure 1c**), which was randomly inserted into the genome. A mosquito line with a single insertion that was mapped to a non-coding locus on the 3^rd^ chromosome was selected (**Figure 1c** and **Supplementary figure 1b,c**; see methods for details). This line, named Fudgel, was further characterized.

**Figure 1.**
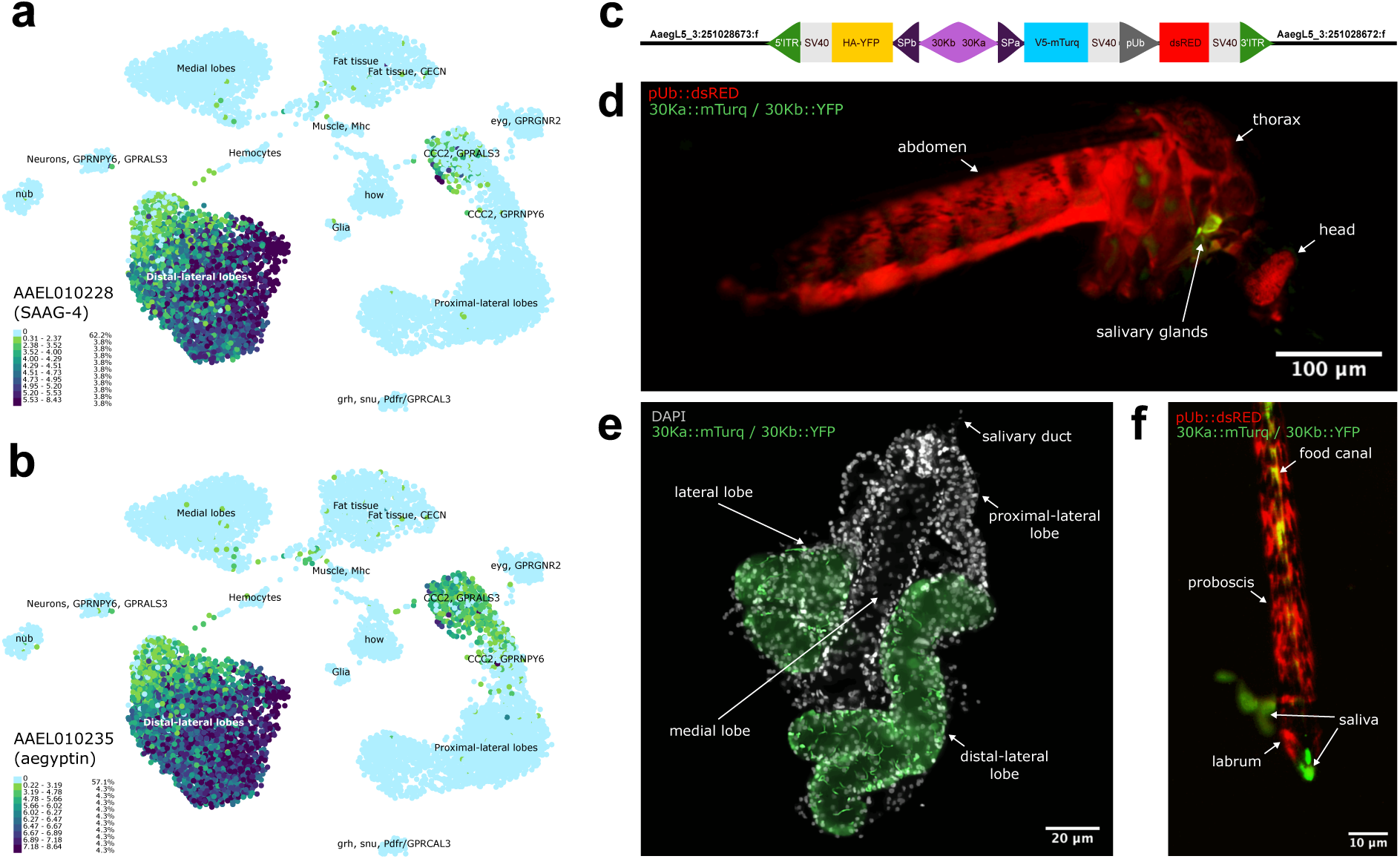
Characterization of the Fudgel transgenic mosquito line to track salivation. **a-b**) UMAP representation of *30Ka* (*AAEL010228*/SAAG-4) (**a**) and *30Kb* (*AAEL010235*/aegyptin) (**b**) expression patterns in *Ae. aegypti* snRNAseq data from salivary glands generated with the UCSC Cell Browser^30^. **c**) Fudgel transgenic construct and insertion locus in the *Ae. aegypti* Bangkok genome. The construct is composed of the bidirectional *30K* promoter (*30Ka* and *30Kb*) and their associated endogenous signal peptides (purple and dark purple). *30Ka* drives the expression of the *mTurq* gene tagged in 5’ with a V5 tag and followed by a SV40 transcription terminator (blue and grey). *30Kb* drives the expression of the *YFP* gene tagged in 5’ with a HA tag and followed by SV40 (yellow and grey). DsRed2 is used as a transgenesis reporter and expressed ubiquitously with a *PUb* promoter and controlled by SV40 (dark grey, red and grey). The construct is flanked by Inverted Terminal Repeats (ITRs) derived from the *piggyBac* transposons and used for transgenesis^35^. The insertion locus of the transgene was identified by Inverse PCR. **d-f**) Expression patterns of the mTurq/YFP (green) and DsRed2 (red) fluorescent markers in whole body (**d**), salivary gland (**e**) and proboscis (**f**) of Fudgel mosquitoes. Nuclei stained with DAPI are shown in **e**.

### Characterizing mosquito salivation using the Fudgel reporter line

The analysis of Fudgel mosquitoes on a fluorescent microscope with a green filter - which does not differentiate YFP and mTurq - revealed fluorescent labeled tissue in the mosquito thorax, where the salivary glands are expected to be (**Figure 1d**). In contrast, the DsRed2 fluorescent marker, whose expression is driven by a ubiquitous promoter, labeled the whole mosquitoes (**Figure 1d**). In dissected salivary glands from Fudgel mosquitoes, green labelling was observed in the distal-lateral lobes (**Figure 1e**) as expected using the 30k promoter. Of note, YFP had a broader expression pattern confirming previous observations that *30Kb* is also expressed in proximal-lateral lobes while *30Ka* had a more restricted expression^28^. On a fluorescent microscope, we noted that the mTurq signal was more intense than YFP. This difference may be due to a genomic position effect of the *piggyBac* insertion, since *30Kb* is normally more expressed than *30Ka*^30^, and aegyptin is more abundant in saliva than SAAG-4^14^. Nevertheless, the expression pattern of the fluorescent proteins in the salivary gland was as expected based on the promoters we used. Since the signal peptides encoded by *30Ka* and *30Kb* are retained in the construct, we next analyzed the presence of the fluorescent markers in the saliva itself. Fluorescent saliva could be observed in green, blue and yellow channels inside the food canal of the proboscis while the walls of the canal are labelled with DsRed2 (**Figure 1f** and **Supplementary Figure 1d**). A real-time video shows release of the mTurq fluorescent marker during salivation on a Petri dish (**Supplementary video 1**). These results suggest that Fudgel mosquitoes produce and secrete fluorescent markers in in the saliva. Although the markers are specifically produced by the distal-lateral lobes, proteins from all lobes are represented in the saliva (**Supplementary table 2**)^14,30^. Therefore, the release of the fluorescent markers should allow direct tracking of salivation.

To quantitatively monitor the release of saliva proteins by individual mosquitoes, we measured levels of fluorescent markers by western blot using a forced salivation protocol (**Figure 2a**). V5-mTurq could be readily detected after 30 minutes of forced salivation in about half of the mosquitoes tested (**Figure 2b,c**). Neither the marker levels nor the proportion of individuals with detectable markers - together referred to as salivation performance - changed significantly at 0.5, 1 and 2 hours of forced salivation (**Figure 2c**). Of note, HA-YFP was not detected by western blot (**Figure 2b**) consistent with the microscopy results that suggested it is not as abundant as V5-mTurq. Using V5-mTurq as a reporter for the amount of saliva proteins, we next analyzed the salivation performance of individual mosquitoes at different stages of completion of their gonotrophic cycle. Blood-fed Fudgel mosquitoes were analyzed at different days post-feeding (dpf) (7 dpf, 10 dpf or 14 dpf) with or without egg laying compared to sugar-fed individuals (**Figure 2d**). Overall, the physiological state of the mosquito did not affect salivation performance (**Figure 2e**). There was a significant decrease in saliva levels at 10 dpf in mosquitoes that were allowed to lay eggs and thus completed their gonotrophic cycle (**Figure 2e**). This tendency at day 10 was observed in three out of four independent replicates indicating a physiological difference rather than experimental variation (**Supplementary figure 4**). This observation is in agreement with previous reported variations in host seeking and blood feeding behavior at different stages of the gonotrophic cycle^36^. We also quantified levels of the fluorescent marker remaining in the salivary glands after salivation (**Figure 2f**). No significant differences in markers levels - mTurq and YFP - were observed in salivary glands, compared to control mosquitoes after blood-feeding on mice or forced salivation (**Figure 2g**). These results suggest that levels of saliva proteins released in both natural and artificial salivation settings are relatively small compared to what remains in the glands.

**Figure 2.**
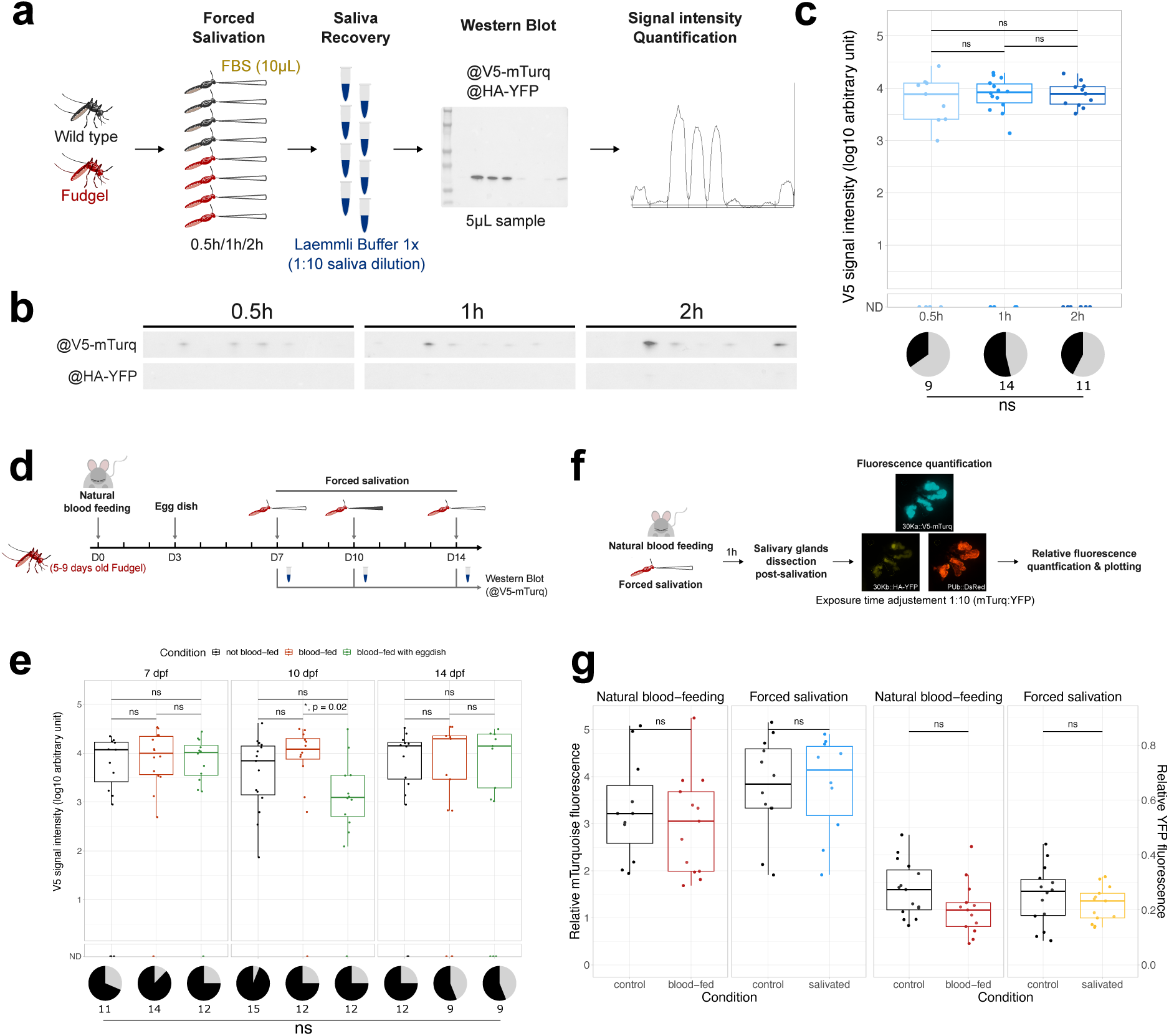
Fluorescence marker detection can be used as a reporter for salivation performance. **a**) Experimental design for detection and quantification of marker expression after forced salivation. **b**) Western Blot signal for V5 and HA in individual saliva samples of Fudgel mosquitoes after 0.5h, 1h or 2h of forced salivation. **c**) V5 signal intensity in individual saliva samples of Fudgel mosquitoes at different times (n = 20 saliva samples per time point) (log10 scale). Pooled results from independent replicates (N = 3). Individual western Blot results are shown in **Supplementary figure 2** and the raw data in **Supplementary table 3**. **d**) Experimental design for testing the effect of different physiological states on salivation. **e**) V5 signal intensity in individual saliva samples of Fudgel mosquitoes in different physiological states after 1h of forced salivation (n = 16 per condition) (log10 scale). Pooled results from independent experiments (N = 4). Individual western Blot results are shown **Supplementary figure 3**, individual plots in **Supplementary figure 4** and the raw data in **Supplementary table 4**. **f**) Experimental design for testing the effect of salivation on salivary glands. **g**) mTurq and YFP fluorescence intensity relative to DsRed2 (CTCF(marker/DsRed2)) in individual salivary glands of Fudgel mosquitoes 1h after blood-feeding on mice or forced salivation. Exposure time was 10-fold higher for YFP compared to mTurq. Representative images are shown in **Supplementary figure 5** and the raw data in **Supplementary table 5**. Comparison of signal intensity: Kruskal-Wallis test, threshold 0.05. Comparison of prevalence of salivation: Fisher’s exact test, threshold 0.05.

The interaction between mosquito saliva proteins and skin cells after biting has been thoroughly studied *in vitro* using mammalian cell lines or recombinant proteins^37–39^. However, *in vivo* analyses are harder to perform due to the lack of tools to precisely identify and isolate mosquito biting sites and track saliva proteins. Tracking mosquito salivation on a mammalian host after biting would allow precise monitoring of viral replication sites and local immune responses. Since we observed active release of mTurq during forced salivation on a Petri dish (**Supplementary video 1**), we next tested whether the fluorescent markers could be detected directly in human skin explants exposed to mosquitoes (**Figure 3a**). Mosquitoes actively probed the skin explants but could not feed on the human skin biopsies likely due to blood coagulation and/or blood vessel collapse within the 3 hours between surgery and the experiment (**Figure 3b**). Fluorescence microscopy allowed visualizing mosquito bite sites via tracking mTurq and YFP fluorescence markers immediately after probing (**Figure 3c**). Of note, DsRed2 expressed under a ubiquitous promoter and lacking a signal peptide was not detected at the bite site (**Figure 3c**). This suggests specific secretion and release of the fluorescent markers expressed under the control of the bidirectional *30K* promoter during probing on the skin explants. Thus, the Fudgel line allows efficient tracking of bite sites and the release of saliva reporters in the skin.

**Figure 3.**
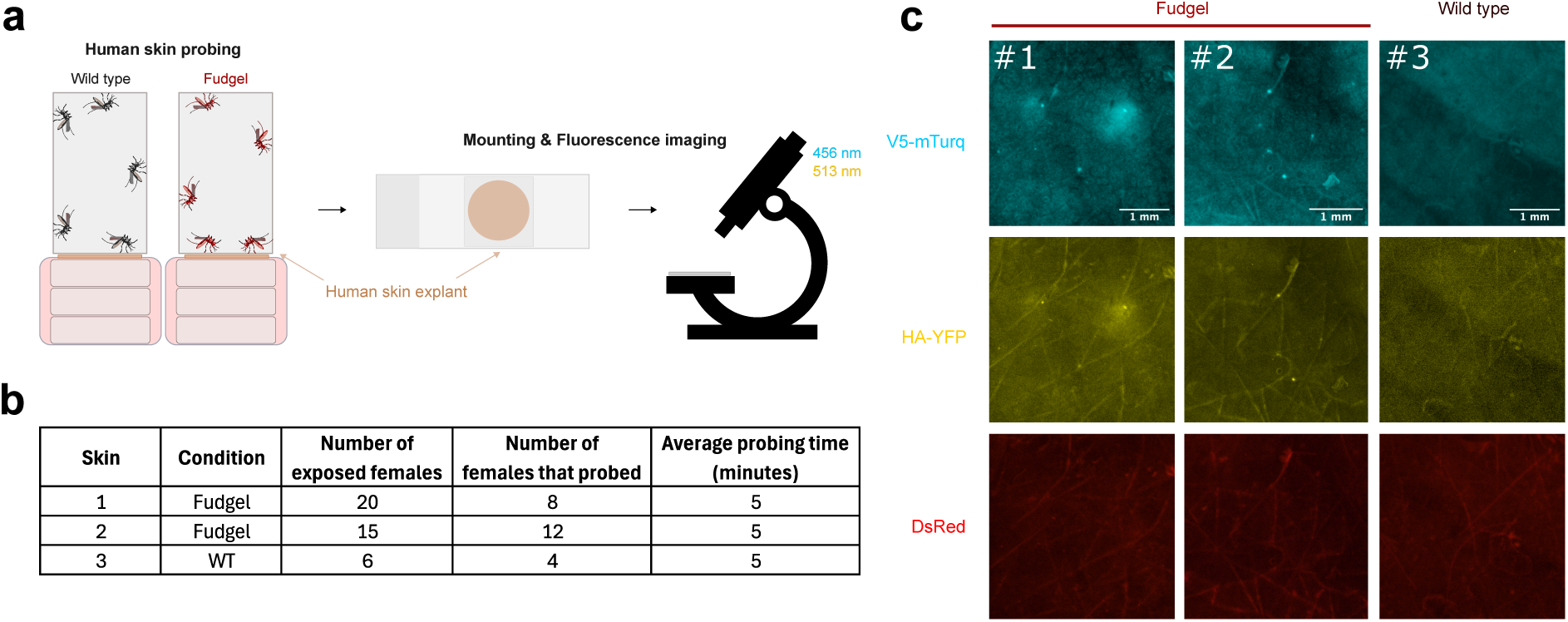
Fluorescent markers allow tracking of probing sites on human skin. **a**) Experimental design for the analysis of probing on human skin explants. **b**) Mosquito exposure condition and time for each skin explant. **c**) Detection of mTurq and YFP fluorescence at probing foci on each tested human skin explant directly after probing. Exposure time was 10-fold higher for YFP compared to mTurq.

### Virus transmission and mosquito salivation

Mosquito-borne viruses are transmitted by mosquitoes through biting and virus detection in forced salivation assays is often used as a measure of transmission^26,27^. However, in these assays, quantifying the release of saliva proteins is rarely performed due to the lack of experimental tools. Mosquito engorgement with buffer or measurements of the volume of release liquid in oil-filled capillary tubes have both been used as a proxy for salivation^22,23^. To overcome these limitations, we used the Fudgel line to investigate the connection between salivation and virus transmission (**Figure 4a**). In the following assays, mosquitoes were infected with ZIKV by intrathoracic injection to guarantee homogenous systemic dissemination, since we wanted to analyze salivary gland infection and virus presence in the saliva. Viral loads in infected Fudgel mosquito carcasses were not significantly different from non-transgenic controls showing that the transgene does not affect infection (**Figure 4b**). Using a forced salivation protocol, we observed that salivation performance was significantly reduced in ZIKV infected Fudgel mosquitoes, which displayed a lower proportion of individuals with detectable saliva reporter in forced salivation samples (26 out of 40) compared to non-infected mosquitoes (16 out of 16) (**Figure 4c)**. mTurq RNA levels in infected individuals were not significantly different from non-infected controls, suggesting that the effect observed in response to ZIKV infection is due to a decrease in protein levels or release of the saliva reporter (**Figure 4d**). We next measured the presence of infectious virus in the saliva samples from the same individual mosquitoes. Out of 40 tested saliva samples from infected Fudgel mosquitoes, 9 had a cytopathic effect on cells (shown in green on **Figure 4c**). In these samples, we observed no correlation between salivation performance and detection of the virus (**Figure 4c**). Of note, two ZIKV-positive samples had no detectable saliva reporter (**Figure 4c**). To verify if the effect of virus infection on salivation was restricted to ZIKV, we next analyzed mosquitoes infected with CHIKV by intrathoracic injection. Fudgel mosquitoes showed similar CHIKV levels compared to controls, showing no effects of the transgene on viral infection (**Figure 4e**). Similar to ZIKV, we observed a decrease in salivation performance in CHIKV infected mosquitoes with a decreased proportion of salivating individuals (38 out of 54) compared to non-infected controls (19 out of 20), and an additional significant decrease in the amount of detected saliva reporter (**Figure 4f**). Similar to ZIKV, *mTurq* RNA levels in CHIKV infected individuals were not significantly different from non-infected controls (**Figure 4g**). Out of 54 samples, 5 were infectious on mammalian cells (**Figure 4f**) and again we observed no correlation between virus detection and the amount of detected saliva reporters.

**Figure 4.**
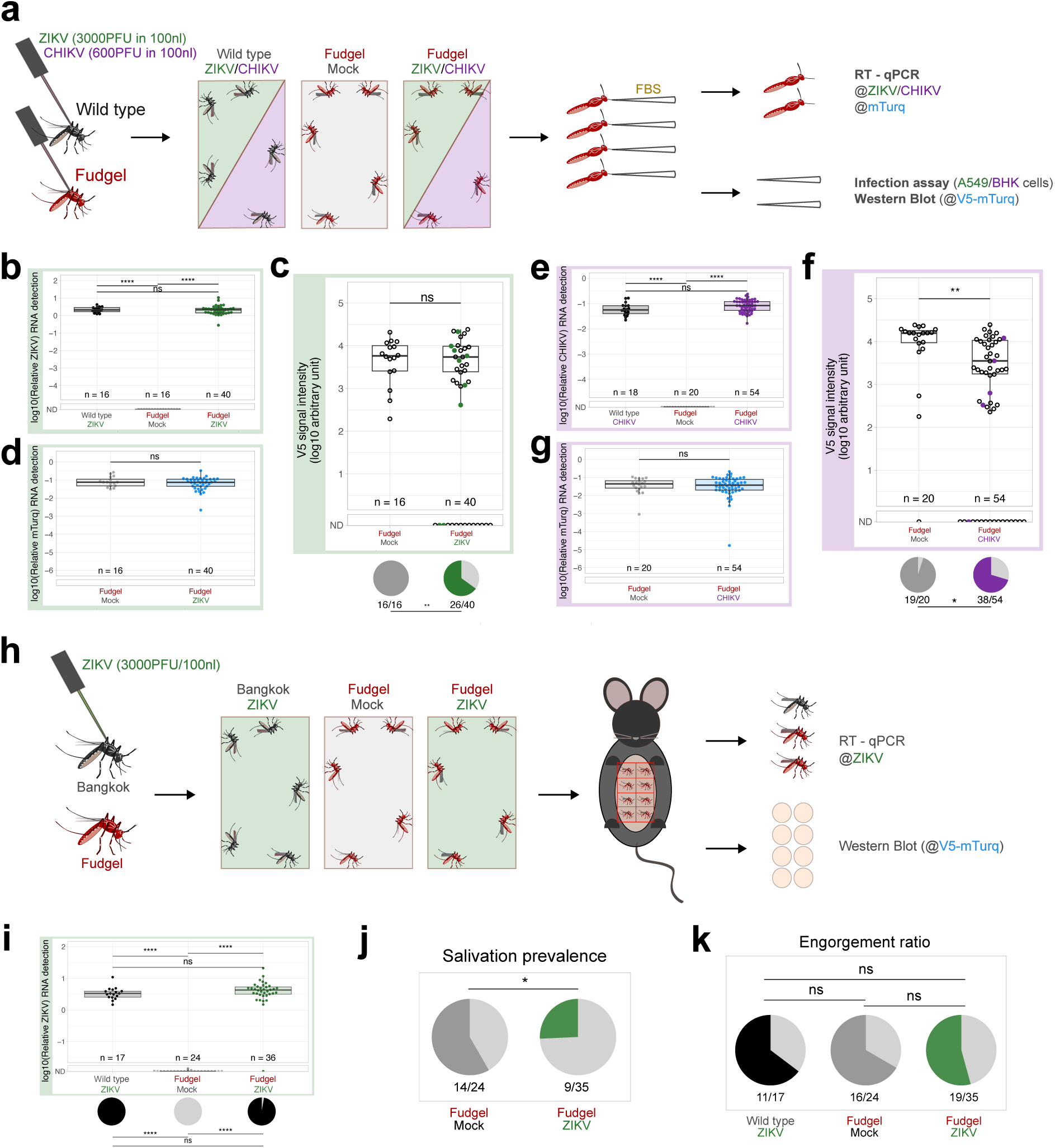
ZIKV and CHIKV decrease salivation in infected mosquitoes. **a**) Experimental design for the analysis of salivation in mosquitoes infected with ZIKV or CHIKV. **b,e**) Detection of ZIKV (**b**) and CHIKV (**e**) by RT-qPCR in mosquitoes at respectively 14 days and 9 days post infection. **c,f**) V5 signal intensity and salivation prevalence in individual saliva samples of control or ZIKV-infected (**c**) and CHIKV-infected (**f**) mosquitoes. Samples associated with the presence of infectious virus are marked respectively in green and in purple. **d,g**) Detection of V5-mTurq expression by RT-qPCR in control and ZIKV-infected (**d**) or CHIKV-infected (**g**) mosquitoes. **b-f** show pooled results from independent experiments (ZIKV N = 2, CHIKV N = 3). Individual Western Blots are shown in **Supplementary figure 6**, individual plots in **Supplementary figure 7** and the raw data in **Supplementary tables 6** and **7**. **h**) Experimental design for the analysis of mice bitten by mosquitoes infected with ZIKV. **i**) Detection of ZIKV by RT-qPCR in mosquitoes at 15- or 16-days post infection. **j**) Prevalence of individual mouse skin samples displaying V5-mTurq signal by western blot after being bitten by a control or a ZIKV-infected mosquito. Pooled results from independent experiments (N = 2). Individual western Blots are shown in **Supplementary figure 8**. **k**) Engorgement ratio of mosquitoes after 10 minutes of contact with the mouse. Comparison of viral loads in carcasses: Kruskal-Wallis test, threshold 0.05. Comparison of V5 signal intensity and relative mTurq RNA level: Student test, threshold 0.05. Comparison of prevalences and engorgement ratios: Fisher’s exact test, threshold 0.05.

We next analyzed the effect of virus infection on salivation performance in a mouse model, which is more physiologically relevant. Mosquitoes were given the opportunity to bite naive mice and the release of saliva was directly measured in mouse skin biopsies (**Figure 4h**). Viral loads in infected Fudgel mosquito carcasses were not significantly different from non-transgenic controls after intrathoracic injection of ZIKV (**Figure 4i**). To monitor the release of saliva reporters, we measured protein levels of fluorescent markers in skin biopsies bitten by individual mosquitoes (**Figure 4h**). In this natural salivation model, a decreased proportion of individuals in ZIKV-infected mosquitoes (9 out of 35) had detectable marker release in skin biopsies compared to non-infected controls (14 out of 24) (**Figure 4j**). Of note, engorgement ratios were not significantly different between infected and non-infected mosquitoes, showing that the observed decrease in salivation performance is not due to a modification of the feeding capacity (**Figure 4k**). Together, these results confirm that virus infection decreases the release of saliva reporters by infected salivary glands.

### Virus infection disrupts the organization of the saliva reporter in the salivary glands

Previous studies have shown changes in the morphology of salivary glands upon infection with CHIKV^11^. To analyze the effect of virus infection on the salivary glands, we analyzed the organization of saliva reporters as well as the segregation of viral antigens by immunofluorescence. Here, we used whole Fudgel mosquitoes embedded in Optimal Cutting Temperature (OCT) compound prior to frozen sectioning on a microtome-cryostat to preserve the organization of salivary glands as much as possible (**Figure 5a**). Whole mosquito sections were analyzed at 15 days post intrathoracic injection of ZIKV to ensure infection of salivary glands. In whole mosquito images, mTurq signal was restricted to salivary glands while ZIKV staining was detected in different parts of the body (**Supplementary figure 9**). In salivary glands, we observed light diffused signal for the saliva reporter that seemed to correspond to the apical cavity of the secretory acinar cells, where saliva proteins are stored (**Figure 5b**). In addition, we observed stronger signal outside of the apical cavity, suggesting accumulation of the reporter in the cytoplasm of acinar cells (**Figure 5b**). This stronger signal of the reporter formed filament-like structures on the lateral and basal sides of the acinar cells that seemed to connect with the salivary duct at the center of the salivary gland (**Figure 5b**). A similar pattern of diffused signal for the saliva reporter in acinar cells was observed in infected salivary glands although the filament-like structures seemed reduced (**Figure 5b**). To quantitate these differences, we used an analysis tool developed for the quantification of mitochondrial filaments using deconvoluted images (**Figure 5c**). This analysis highlighted a significant decrease in the mean area (µm^2^) of the saliva reporter filament-like structures in ZIKV-infected compared to non-infected salivary glands (**Figure 5d**). These results show a disruption of the segregation pattern of the saliva reporter in the acinar cells caused by the virus. Furthermore, the loss of mTurq clusters seemed to occur in areas where there was accumulation of the viral envelope protein (Supplementary **Figure 10a**). To investigate this hypothesis, we further analyzed salivary glands displaying the strongest staining for the envelope protein of ZIKV (**Figure 5e** and **Supplementary Figure 10b**). Using these images, we calculated Manders’ overlap coefficients (M1 and M2) to quantify the colocalization between mTurq and ZIKV. We observed a small fraction of the signal for the ZIKV envelope protein (<0.1) had any overlap with the mTurq filaments, which suggests segregation of the two proteins (**Figure 5f**). A small but more significant fraction of the mTurq filaments (<0.3) overlapped with the signal for the ZIKV envelope (**Figure 5f**). Given that mTurq filaments are uniformly present throughout uninfected salivary glands (**Supplementary Figure 10a),** we may infer that the buildup of viral antigens during replication in acinar cells leads to the displacement of saliva proteins. That is why the overlap of the ZIKV signal is always low because it only reflects the origin. In contrast, the mTurq signal is everywhere at the start of the infection, which means there is always some overlap with ZIKV antigens.

**Figure 5.**
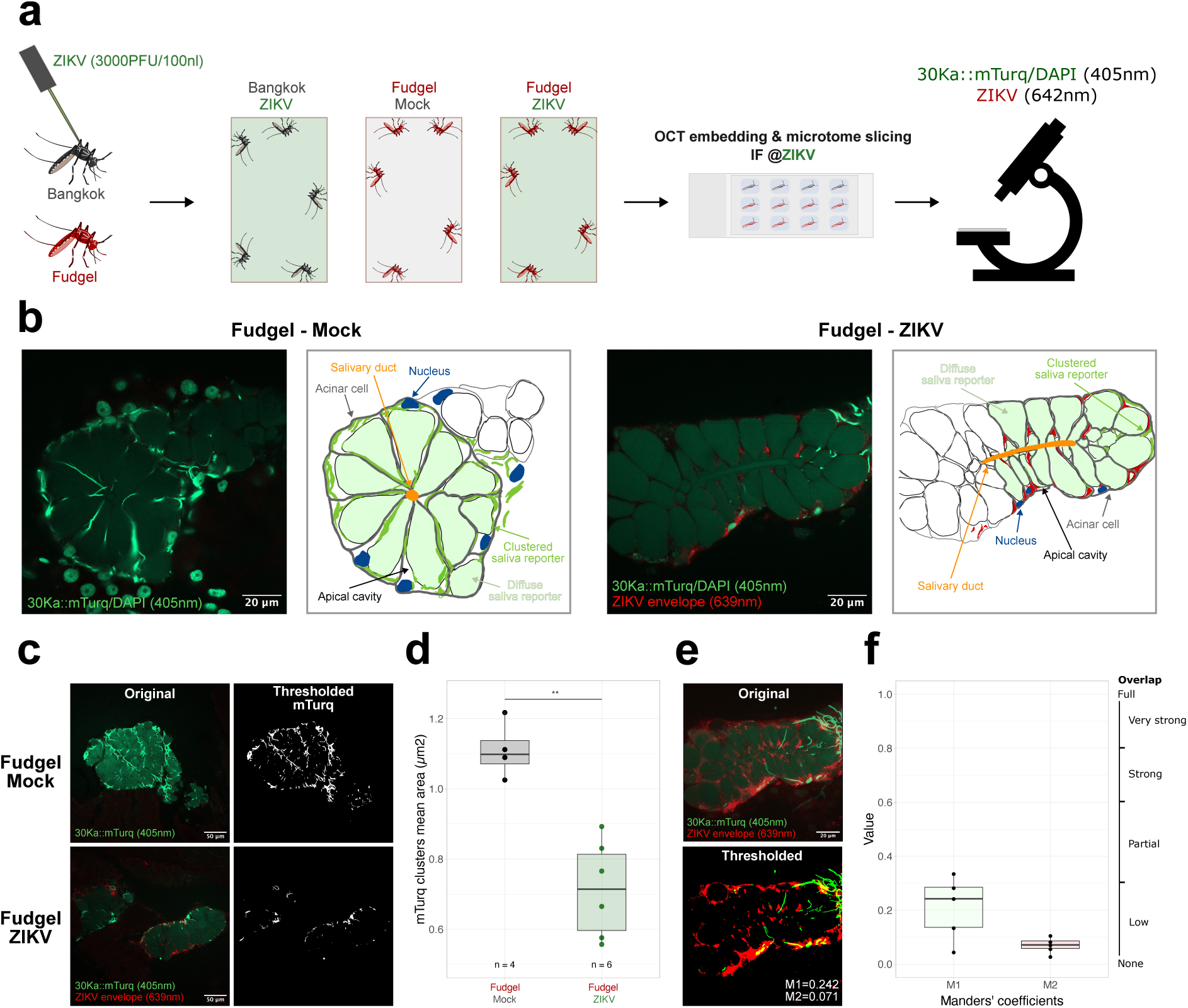
ZIKV infection affects the organization of saliva reporters in infected salivary glands. **a**) Experimental design for immuno-fluorescence analysis of ZIKV-infected mosquitoes. **b**) Representative confocal images of a mock (transverse section) and a ZIKV-infected salivary gland (longitudinal section). A representative scheme highlighting acinar cells (grey), salivary duct (orange), nuclei (blue), saliva reporter (green) and ZIKV envelope (red) is shown on the right. **c**) Representative confocal images of Mock and ZIKV-infected salivary glands showing staining for the saliva reporter (mTurq in green) and the ZIKV E protein (in red). Confocal images (“Original”) were deconvoluted using the “Iterative Deconvolve 3D” plugin^40^ in ImageJ (20 iterations) and saliva aggregates were highlighted using the “2D Threshold” tool on the “Mitochondria Analyzer” plugin^41^ (“Thresholded”). **d**) Mean area (µm^2^) comparison between saliva reporter filaments in Mock and ZIKV-infected salivary glands. Wilcoxon test, threshold 0.05. Mock (n = 4) and ZIKV-infected (n = 6). Images of individual salivary glands are shown in **Supplementary figure 10a.** Calculated area values for each individual salivary gland are in **Supplementary table 9**. **e**) Representative confocal image, original and deconvoluted, showing infected salivary glands from Fudgel mosquitoes stained for the salivary gland reporter and the ZIKV envelope protein. **f**) Manders’ overlap coefficients indicating the fraction of mTurq that overlaps with ZIKV E staining (M1) and the fraction of ZIKV E that overlaps with mTurq (M2). Individual images and calculations are shown in **Supplementary figure 10b**.

## Discussion

In this study, we developed a transgenic mosquito line expressing fluorescent markers that function as reporters for saliva proteins. This line allowed us to study basic aspects of the physiology of salivation. We first analyzed the evolution of salivation behavior through a gonotrophic cycle, e. g. from eggs development to their oviposition. We could highlight a transient decrease in salivation performance at 10 dpf for mosquitoes completing of a full cycle, which was observed neither in mosquitoes that were not allowed to lay eggs after feeding nor in those who did not blood feed. Previous studies suggest hormonal variations through the reproduction cycle of female mosquitoes which would be associated with modifications in host seeking and blood feeding behaviors^36^. The observed transient decrease in salivation performance goes along with a decreased host seeking behavior associated with the start of a new gonotrophic cycle. This phenomenon would need to be further characterized, and our tool is a great asset for this purpose. We also measured the effect of natural blood-feeding or forced salivation on salivary glands purging, without detecting any significant difference in saliva content compared to not blood-fed or non-salivating controls. Moreover, this tool allowed us to precisely visualize mosquito probing sites on human skin explants, highlighting its potential use for *in vivo* transmission studies. This could be an important tool to examine skin cells response to saliva components specifically at the biting site, in both non-infected and virus-infected settings. Overall, our results illustrate the potential of the Fudgel reporter line to help gain valuable insights on the physiology of mosquito salivation.

In the context of infection, we showed that both ZIKV or CHIKV decreased mosquito salivation, strongly decreasing the proportion of mosquitoes that secreted detectable levels of saliva reporters. Notably, virus infection disrupted the organization and decreased levels of the saliva reporters in the secretory acinar cells of salivary glands. Of note, viruses did not affect the RNA levels of markers in the salivary gland, thus suggesting a specific effect of these viruses at the protein level. Indeed, others have shown that expression and secretion of saliva proteins can be impacted by virus infection in mosquitoes^42,43^. In addition, our results show that viral antigens are segregated from saliva proteins in infected acinar cells. These observations are in agreement with previous electron microscopy studies of CHIKV-infected salivary glands which suggested that viral particles segregate in separate vesicles from saliva proteins in acinar cells^11^. We propose that as the viral factory is assembled and antigens accumulate, this displaces saliva proteins. Thus, we observe overlap between viral antigens and saliva reporters only at the borders where viral factories are initiating. Then, as the virus replicates and antigens accumulate, saliva proteins are completely gone. As a result of this segregation from saliva proteins, viral particles could presumably be secreted independently. This provides a mechanistic explanation for the disconnection between release of saliva proteins and viral transmission. Importantly, we observed no major damage to acinar cells, this suggesting the cells remain viable, which may be important to allow efficient and continuous virus replication. Our results are in agreement with previous observations that virus titers do not correlate with the volume of saliva released in forced salivation assays^25^.

Our work represents a shift from the common assumption that viral transmission efficiency correlates with the release of saliva proteins. Of note, genetic depletion of salivary glands or microsurgery removal of the salivary duct caused a reduction in salivation and increased probing behavior in *Anopheles* mosquitoes^44,45^. Thus, reduced release of saliva in virus-infected mosquitoes should lead to an increase in probing and biting behavior. Accordingly, previous studies have shown that dengue-infected mosquitoes have increased biting behavior compared to uninfected controls ^46^. This makes sense in light of data showing that successive short probing events resulted in multiple deposition of the virus^46^. Damaging the salivary glands could thus be a strategy used by the virus to reduce salivation and increase the probing behavior to enhance viral transmission. A similar situation might occur in *Anopheles* mosquitoes, where there are significant changes in salivary glands morphology and loss of integrity upon *Plasmodium* infection^47^. The authors observed an alteration of saliva composition with a decrease of detected proteins and diffusion of saliva proteins in the mosquito hemolymph. At the same time, no difference was observed in the volume of saliva released from *Plasmodium*-infected mosquitoes although they seemed to probe for longer times^44,47^. Overall, these data suggest that increased probing would serve to increase transmission of pathogens by the mosquito bite. Our results may also help explain recent epidemiological observations that low levels of circulating antibodies against the D7 *Aedes* salivary protein correlate with a higher infection burden of DENV in human populations living in *Aedes* endemic populations^48^. Hence, we propose here a transmission model in which viral infection reduces the secretion of saliva proteins by the mosquito, resulting in increased probing and enhanced delivery of virus particles independently of saliva. Given the disconnect between salivation and release of viral particles, it will be important to further investigate unclear how the virus is delivered during mosquito probing. We believe that the tool we developed here will help answer these questions and help develop strategies to control the transmission of vector borne viruses.

## Material and methods

### Cloning, transgenesis and Fudgel line establishment

Sequences of the 30K bidirectional promoter and associated signal peptides were recovered from the AaegL5 LVP_AGWG reference genome on the VectorBase database (https://vectorbase.org/vectorbase/app) and synthesized as a gBlock gene fragment (IDT). The gBlock sequence was cloned using the NEBuilder® HiFi DNA Assembly Cloning kit (ref E5520S) in a pSKB-plasmid (Addgene #62540), previously linearized by PCR amplification using the ThermoFisher Scientific DreamTaq DNA polymerase (ref EP0701) (primers in **Supplementary table 8**). The sequences for epitope-tagged fluorescent markers were amplified from lab stock plasmids by PCR using the ThermoFisher Scientific Phusion High-Fidelity DNA polymerase (ref F530S) (primers in **Supplementary table 8**). A backbone plasmid containing the PUb::DsRed2::SV40 sequence was digested using BamHI and SmaI restriction enzymes (ThermoFisher). The final construct was assembled using NEBuilder® HiFi DNA Assembly Cloning kit (ref E5520S). Full plasmid sequence is provided in **Supplementary Table 8**.

This plasmid was injected at a concentration of 300 ng/µL in mosquito eggs with 100 ng/µL of a helper, codon-optimized *piggyBac* transposase-expressing plasmid^35^ (available on request) in 0.5x PBS as follows: *Ae. aegypti* female mosquitoes of the Bangkok strain were blood-fed 4 days prior to injection and allowed to lay eggs. One hour-old eggs were aligned along wet nitrocellulose on a microscope slide and injected in their posterior pole using a Nikon ECLIPSE TE2000-S inverted microscope, Eppendorf Femtojet 4x micro-injector and Eppendorf TransferMan NK2 control panel. Injected G0 larvae were screened and individuals displaying transient DsRed2 fluorescence expression were backcrossed to wild-type Bangkok mosquitoes. From their G1 progeny, DsRed2-positive individuals were selected under a fluorescent binocular microscope (Nikon SMZ18 microscope, sola light engine Lumencor® light source), backcrossed, and progeny was again individually backcrossed. In G3, populations showing 50% inheritance of the transgene were selected, suggestive of a single transgene insertion. Positive G3 mosquitoes of a given population were self-crossed. Homozygous – single insertion – G4 mosquitoes were sorted based on fluorescence intensity using the COPAS instrument^49^ to generate the Fudgel line (**Supplementary figure 1b,c**). Experiments were carried on using mosquitoes from G5 on.

### Transgene insertion mapping by inverse PCR

Genomic Fudgel DNA was extracted from five homozygous male pupae using the Monarch DNA purification kit (New England Biolabs ref T3010S/L) and quantified with Nanodrop. Extracted DNA (1µg) was digested with AarI, AgeI, ApeI, SacII and XhoI restriction enzymes (FastDigest — Thermo Scientific) for 1 hour at 37°C and enzymes were inactivated for 10 minutes at 80°C. The digested DNA was diluted and treated with T4 DNA ligase (50 Units of enzyme for a 500µL reaction, Thermo Scientific ref EL0011) overnight at room temperature. DNA was then purified using the Macherey-Nagel NucleoSpin Gel and PCR Clean-up kit (ref 11992242) and amplified by nested-PCR using the ThermoFisher Scientific DreamTaq DNA polymerase (ref EP0701). Primers for both PCR reations were designed to amplify the 3’ region of the transgene (**Supplementary table 8**). 15µL of the PCR were Sanger sequenced (GATC Biotech, Eurofins) and the obtained sequence blasted on the AaegL5 LVP_AGWG reference genome on VectorBase. This alignment returned 1 hit on chromosome 3 (AaegL5_3:251028673:f). Deduced flanking genomic sequences were confirmed on both the 5’ and 3’ sides of the *piggyBac* element by PCR amplification using the ThermoFisher Scientific Dreamtaq Green MasterMix (ref K1081) (primers in **Supplementary table 8**) and sequenced (GATC Biotech, Eurofins).

### Salivation protocol optimization

Fudgel female mosquitoes were starved 24h prior to experiment, then anesthetized with CO_2_ and kept on ice. Legs and wings were removed, and proboscis inserted in tips containing 10µL of FBS (Fetal Bovine Serum). Mosquitoes were left to salivate either 30 minutes, 1 hour or 2 hours, then saliva samples were collected. Each mosquito was considered a biological replicate.

### Marker detection and quantification by Western Blot

Saliva samples were diluted 1:10 in 1x Laemmli Buffer and heated for 20 minutes at 95°C. Heating process was repeated depending on experimental set up. Samples were then resolved on Mini-PROTEAN TGX Stain-Free Precast Gel 4-15% (Bio Rad ref #4568086) and transferred to a Nitrocellulose membrane (Trans-Blot Turbo Transfer Pack Bio Rad ref #1704158). Membranes were blocked with 5% BSA (Albumine, from bovine serum SIGMA ref A3059) in PBS-Tween 0.1% solutions and treated with either @HA-HRP (Roche ref 12013819001) or @V5-HRP (Invitrogen ref 46-0708) antibodies 1:5000 in 1.5% BSA in PBS-Tween 0.1%. Membranes were incubated in the antibody solution under agitation either 1 hour at room temperature or overnight at 4°C, rinsed in PBS-Tween 0.1%. Protein signal was revealed with ECL Select (Cytiva ref GERPN2235) using a ChemiDoc imaging system (BioRad). Western Blot images were treated with the ImageJ software. Rectangular selection of signal from all samples of the membrane constitutes a “lane”: *Analyze/Gels/Select First Lane*. Bands intensity was then plotted: *Analyze/Gels/Plot Lanes*, and relative band intensities recovered with the *Wand* tool. Intensity values were then recorded, plotted on a log10 scale, and processed with the RStudio software with the *ggplot2* and associated packages^50,51^. Saliva tag signal levels between conditions were compared using a Kruskal-Wallis test.

### Fluorescent marker imaging

Mosquitoes were anaesthetized with CO_2_ and kept on ice. Whole body and proboscis images were acquired on a Nikon SMZ18 fluorescent binocular microscope with sola light engine Lumencor® (**Figure 1d,f**). For salivary gland acquisitions (**Figure 1e**), dissections were performed in PBS under a fluorescence microscope and the salivary glands were fixed in PFA 4% for 1h40 at room temperature (RT). Samples were then permeabilized in PBS-T (0.2% Triton X-100, 1% BSA) for 45 minutes at RT and incubated with DAPI (1:1000 in PBS-T) for 10 minutes at RT. Salivary glands were rinsed 3 x 10 minutes in PBS-T and mounted on slides using VECTASHIELD Antifade mounting medium (Vectorlabs, ref H-1000). Acquisitions were performed on an epifluorescence microscope Axioskop 2 (Zeiss). For confocal image acquisitions (**Supplementary figure 1d**) salivary glands and mosquito heads were dissected in PBS under a fluorescent stereoscope. Fresh tissues were mounted in VECTASHIELD mounting medium and confocal 10x Z stack acquisitions were performed using a spinning disk inverted microscope (Zeiss, Axio Observer Z1). Stack images were merged using the “Extended Depth of Field” plugin in ImageJ ^52^: Direct Method, Expert Mode, EDF “Complex Wavelets”. For proboscis, acquired images were despeckled (*Process/Noise/Despeckle*) prior to merging to avoid signal loss.

### Salivation recording

Mosquitoes were anaesthetized with CO_2_ and kept on ice. Legs and wings were removed, and proboscis inserted in FBS drops. Mosquitoes were left to salivate at room temperature for 1 hour. Mosquitoes and droplets were then observed under a fluorescent microscope (Zeiss SteREO Discovery.V8 microscope, X-Cite Xylis Excelitas Technologies laser) to witness salivation and for video recording (**Supplementary video 1**).

### Physiological state and salivation

5 to 8 days old Fudgel mosquitoes were blood-fed or not on mice anaesthetized with a solution of NaCl 0.9%, Rompun 2%, Zoletil 50. 3 days post-feeding fed females were given or not an egg dish to lay their eggs. Egg dishes were removed after 3 days. Of note, blood-fed females without the egg dishes had minor residual oviposition on the sugar pads. At day 7, day 10 and day 14 post-feeding, mosquitoes were anaesthetized with CO_2_, legs and wings removed, and females from the different feeding conditions were put to salivate in 10µL FBS for 1 hour. 18 hours prior to salivation, mosquitoes were starved by replacing sugar with water only. Non blood-fed Bangkok females of the same age were used as negative control. Individual saliva samples were collected and diluted 1:10 in Laemmli 1x. The samples were then analyzed by Western Blot with @V5-HRP antibody treatment.

### Salivary gland fluorescence intensity after natural blood feeding or forced salivation

For blood feeding, mosquitoes were starved the prior day. Half of mosquitoes were allowed to fully engorge on a mouse anaesthetized with a solution of NaCl 0.9%, Rompun 2%, Zoletil 50. For forced salivation, mosquitoes were anaesthetized with CO_2_ and kept on ice. Legs and wings were removed, and half of them were put to salivate in 10µL FBS for 1 hour at RT. The salivary glands of control mosquitoes, blood-fed mosquitoes, and mosquitoes that salivated were dissected under a fluorescent binocular microscope (Nikon SMZ18 microscope, sola light engine Lumencor® laser). For each condition of each replicate, between 6 and 8 salivary glands were dissected and analyzed. Pictures were acquired with a fluorescent binocular microscope (Nikon SMZ18 microscope, sola light engine Lumencor® laser). Fluorescence intensity quantification was done using ImageJ. For mTurquoise, YFP and DsRed2, we calculated the Corrected Total Cell Fluorescence (CTCF):

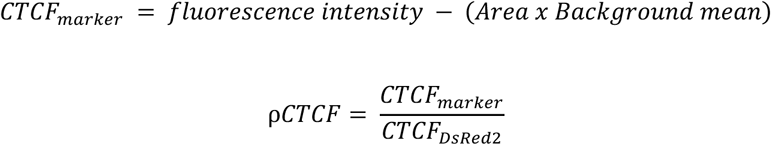

Fluorescence intensity was normalized using ρCTCF, defined as the ratio between the CTCF value of the salivation marker – either mTurq or YFP – and the CTCF of the DsRed2 transgenesis reporter. ρCTCF for each marker of individual salivary glands are plotted using the *ggplot2* and associated packages^50,51^ on RStudio.

### Saliva detection in human skin explants

Human skin explants were recovered from anonymized patients who underwent abdominal reduction (St Anne Clinic, Strasbourg), with written informed consent and institutional review board approval, in agreement with the Helsinki Declaration and French legislation. At reception, skin surfaces were wiped successively with 70% ethanol, antiseptic solution (Dakin®) and phosphate buffer saline (Corning). Superficial layers of the skins (200µm thickness) were harvested using a dermatome blade and incubated in complete DMEM medium (10% fetal calf serum, 0.1mg/mL streptomycin, 100UI/mL penicillin and 10μg/mL gentamycin) overnight at 37°C, 5% CO_2_. The night prior to experiment, 5 female mosquitoes were put in drosophila culture tubes and starved until the experiment. Human skins explants were put in contact with starved mosquitoes and each batch of mosquitoes was allowed to probe for 10 minutes. Different skin samples were then recovered, placed in 1x PBS and immediately observed under a fluorescent binocular microscope (Nikon SMZ18 microscope, sola light engine Lumencor® laser). Pictures were acquired using a Nikon DS-Ri2 camera.

### ZIKV and CHIKV infections

One week old Bangkok and Fudgel female mosquitoes were transferred to a Biosafety Level 3 laboratory, anesthetized with CO_2_ and kept on ice. Injections were performed using a Nanoject III micro-injector (Dummond Scientific Company Cat. 3-000-207). Both ZIKV and CHIKV were tested in independent experiments, in duplicates and triplicates respectively. Bangkok females were injected with either 3000 PFU of ZIKV or 600 PFU of CHIKV in 100nL. Fudgel females were injected with either Mock solution (virus growth medium: Gibco Leibovitz’s L-15 Medium), or with 100nL of virus. They were left to recover and incubate. Forced salivations were performed at either 14 days post-injection (dpi) or 9 dpi respectively for ZIKV and CHIKV.

### Forced salivation assays

Mosquitoes were starved (deprived of sugar and water) the day before the experiment and anesthetized with triethylamine (N,N-Diethylethanamine, SIGMA-Aldrich ref T0886). Legs and wings were removed, and proboscis inserted in tips filled with 10µL FBS. Mosquitoes were left to salivate for 1 hour at RT. Saliva samples and carcasses were then recovered for analysis. ZIKV and CHIKV RNA levels in carcasses were measured by RT-qPCR (primers in **Supplementary table 8**). RNA was extracted using the Mag-Bind Total RNA kit (Omega bio-tek ref M6731-01) at half-volumes and the KingFisher Apex machinery (Thermo Scientific). Reverse transcription was performed using RevertAid H Minus enzyme (Thermo Scientific ref EP0452) with random hexamers, and qPCR with PowerSYBR Green PCR Master Mix (Applied Biosystems ref 4367659) and the reaction ran on Quant Studio 5 machinery (Applied Biosystems). Results were processed using the apps.thermofisher.com platform and plotted with *ggplot2* and associated packages^50,51^ on RStudio. To quantify salivation levels, 2µL of each saliva sample were mixed with 18µL of Laemmli 1x (1:10) and analyzed by Western Blot with @V5-HRP antibody. Signal intensity was quantified using ImageJ and plotted with *ggplot2* and associated packages^50,51^ on RStudio.

Of the remaining saliva samples from CHIKV experiments, 5µL were used to perform an infection assay on BHK cells. Cells were previously grown in DMEM medium (10% FBS, 1% Penicillin/Streptomycin (P/S)) at 37°C, 5% CO2. 24h prior to infection, cells were diluted to 10^-4^ cells/100µL in DMEM medium (5% FBS, 1% P/S) and sampled in 96 well plates. Saliva samples were diluted in series in DMEM medium (0% FBS, 2% antibiotic-antimycotic, 1% P/S) and put on cells, from 10^-2^ to 10^-7^ dilution, 100µL/well, each dilution in quadruplicates. Viral stock dilutions from 10^-4^ to 10^-9^ in quadruplicates were used as a positive control for cells infection. Non-infected saliva samples were used as negative controls. Cells were left to incubate with the virus at 37°C, 5% CO_2_. Cytopathic effect was measured from 3dpi to 7dpi. For ZIKV saliva samples, 2µL of each individual saliva samples were used to infect interferon-deficient A549 cells. Cells were grown and prepared as detailed for BHK cells. Saliva samples were diluted in 100µL of DMEM medium (0% FBS, 2% antibiotic-antimycotic, 1% P/S) and put on cells (50x dilution). Each saliva sample was used on one single cell well. Viral stock dilutions from 10^-4^ to 10^-9^ in triplicates were used as a positive control for cells infection. Non-infected saliva samples were used as negative controls. Cells were left to incubate with the virus at 37°C, 5% CO_2_. Cytopathic effect was measured from 3dpi to 7dpi.

### *In vivo* salivation assays

Mosquitoes were starved (deprived of sugar and water) and B6.Cg-ifngr1tm1Agt ifnar1tm1.2Ees/J mice were shaved on their abdomen and transferred to a Biosafety Level 3 laboratory the day before the experiment. For the feeding experiment, mice were anaesthetized with a solution of NaCl 0.9%, Rompun 2%, Zoletil 50. Mosquitoes – respectively 15 and 16 dpi for each experimental replicate – were anesthetized with CO_2_ and transferred into a 3D-printed individual females feeding system comprising 8 single-mosquito chambers (plans available on demand). Each feeder containing 8 individual female mosquitoes was placed on the abdomen of a mouse and left feeding for ten minutes. After feeding, mosquitoes were anesthetized with CO_2_ and kept on ice. Engorgement status for each female was noted, and carcasses were then recovered for analysis. ZIKV and CHIKV RNA levels in carcasses were measured by RT-qPCR (primers in **Supplementary table 8**). RNA was extracted using the Mag-Bind Total RNA kit (Omega bio-tek ref M6731-01) at half-volumes and the KingFisher Apex machinery (Thermo Scientific). Reverse transcription was performed using RevertAid H Minus enzyme (Thermo Scientific ref EP0452) with random hexamers, and qPCR with PowerSYBR Green PCR Master Mix (Applied Biosystems ref 4367659) and the reaction ran on Quant Studio 5 machinery (Applied Biosystems). Results were processed using the apps.thermofisher.com platform and plotted with *ggplot2* and associated packages^50,51^ on RStudio. Anesthetized mice were euthanized by cervical dislocation and the bite sites on the abdomen skin regions were individually recovered with 8mm disposable biopsy punches (Kai medical ref BP-80F) and put in TRIzol LS Reagent (Invitrogen ref 10296010). Proteins of individual skin samples were extracted using TRIzol Reagent Invitrogen protocol (catalog numbers 15596026 and 15596018), resuspended in 200µL TrisHCl 50mM, SDS 1x, protease inhibitor + EDTA and diluted in 200µL Laemmli buffer 2x (1:2). Individual skin protein samples were analyzed by Western Blot with @V5-HRP antibody and @Actin Monoclonal Antibody (Invitrogen mAbGEa ref MA1-744) 1:5000 in 1.5% BSA in PBS-Tween 0.1%.

### Microtome-slicing and immuno-fluorescence of ZIKV-infected mosquitoes

Mosquitoes were anesthetized with CO_2_ 15 dpi and kept on ice. They were pre-fixed by injection of 100nl PFA 16% and fixed by immersion in PFA 4% for 2h at RT. They were then dehydrated by successive baths in increasing concentration of sucrose solutions at RT: 12 hours in PBS 15% sucrose, and 24h in PBS 30% sucrose. Dehydrated samples were pre-embedded for 48h at RT in liquid OCT compound (Cellpath ref KMA-0100-00A). Embedding was done by immersing samples in liquid OCT compound in plastic molds (Fisher Disposable Base Molds ref 22-363-353) and freezing in liquid nitrogen. Samples were kept at -20°C. OCT blocks were microtome-sliced using a Cryostat system (Leica CM3050 S) and 30µm slices were recovered on Clean SuperFrost slides (Epredia Superfrost Plus Adhesion Microscope Slides ref J1800AMNZ). Slides were allowed to dry 2h at RT, immersed in ice cold acetone for 30 minutes at RT and dried for 30 minutes. Each sample was circled with a hydrophobic barrier pen (ThermoFisher ReadyProbes Hydrophobic Barrier Pap Pen ref R3777). Samples were blocked with PBS-BSA 2% 2 times 10 minutes at RT and permeabilized in PBS-BSA 1% + Triton 0.1% for 45 minutes at RT. Samples were incubated in primary antibody against ZIKV (mouse 4G2, laboratory synthetized stocks) at 1:50 in PBS-Triton 0.1% overnight at 4°C and washed 3 times 10 minutes in PBS-Triton 0.1%. Samples were blocked and permeabilized in PBS-BSA 1% + Triton 0.1% for 45 minutes at RT, and incubated in secondary antibody (Invitrogen Alexa Fluor 647 goat anti-mouse IgG ref A21235) for 2h at RT. They were then washed 2 times 10 minutes in PBS-Triton 0.1% and incubated with DAPI at 1:500 in PBS-Triton 0.1% for 15 minutes at RT. They were washed once in PBS-Triton 0.1%, mounted in VECTASHIELD mounting medium and confocal 20x Z stack acquisitions were performed using a confocal microscope (Zeiss, Beam Path LSM980). Images were processed in ImageJ. 10x images were adjusted using the “Enhance Contrast” tool (saturation 405nm channel 0.05%, 639nm channel 0.1%). The saliva marker patterns were analyzed following a mitochondrial quantification method optimized by Hemel et al.^53^. 40x images were deconvoluted using the “Iterative Deconvolve 3D” plugin in ImageJ^40^ (20 iterations) and saliva aggregates were highlighted using the “2D Threshold” tool on the “Mitochondria Analyzer” plugin^41^. The mean area (µm2) of mTurq aggregates in Mock and ZIKV-infected salivary glands was extracted from the “2D Threshold” analysis, plotted using the ggplot2 package on RStudio and analyzed using a Wilcoxon test, threshold 0.05. 63x stack images were deconvoluted using the “Iterative Deconvolve 3D” plugin in ImageJ^40^ (20 iterations) and saliva aggregates were highlighted using a Otsu’s Thresholding method. Correlation tests were performed on whole stacks using the “JaCoP” plugin^54^. As for representation, stack images were merged using the “3D Project” tool.

## Acknowledgements

We thank members of the Mosquito Immune Response team for comments and discussions and staff at UPR9022 and U1257 for technical support, and especially Marcela Celis for maintenance of mosquito lines. This work was supported by grants from Conselho Nacional de Desenvolvimento Científico e Tecnológico (CNPq) to J.T.M.; Fundação de Amparo a Pesquisa do Estado de Minas Gerais (FAPEMIG), Instituto Nacional de Ciência e Tecnologia de Vacinas (INCTV) to J.T.M.; Fonds régional de coopération pour la recherche FRCT2020 Région Grand-Est – ViroMod to J.T.M. and C. M.; Institute for Advanced Studies of the University of Strasbourg (USIAS fellowship 2019) to J.T.M.; Human Frontiers Since Program (RGP018/2023) to J.T.M.; Agence Nationale de la Recherche ANR-21-CE12-0024 and ANR-24-CE15-4470 to J.T.M. and , ANR-19-CE35-0007 to E.M. and J. T. M.. This study was financed in part by the Coordenação de Aperfeiçoamento de Pessoal de Nível Superior — Brasil (CAPES) — Finance Code 001 to J.T.M. This work of the Interdisciplinary Thematic Institute IMCBio, as part of the ITI 2021-2028 program of the University of Strasbourg, CNRS and Inserm, was supported by IdEx Unistra (ANR-10-IDEX-0002), by SFRI-STRAT’US project (ANR 20-SFRI-0012), and EUR IMCBio (IMCBio ANR-17-EURE-0023) under the framework of the French Investments for the Future Program to J.T.M.. L.S. was supported by fellowships from IMCBio and FRM (Programme Fin de Thèse FDT2024040 18199). H.M. was supported by a FRM Postdoctoral Fellowship (SPF202110013925). The funders had no role in study design, data collection and analysis, decision to publish, or preparation of the manuscript.

## Author contributions

L.S., M.B. and J.T.M. designed the project. L.S., M.B., B.V., T.H.J.F.L., E.M. and F.A. performed experiments. L.S. and J.T.M. analysed the data. L.S. and J.T.M. wrote the paper. All authors discussed the results and edited the manuscript.

## Competing interests

The authors declare no competing interests.

## Supplementary Materials

### Supplementary Figures

**Supplementary figure 1-** Establishment and phenotypic characterization of the Fudgel transgenic line.

**Supplementary figure 2-** Quantification of saliva reporter levels by Western Blot.

**Supplementary figure 3-** Quantification of the saliva reporter in samples collected from mosquitoes in different physiological states.

**Supplementary figure 4-** Individual replicate plots for @V5-mTurq detection in saliva samples collected from mosquitoes in different physiological states.

**Supplementary figure 5-** Fluorescent images of salivary glands from Fudgel mosquitoes after natural blood-feeding or forced salivation.

**Supplementary figure 6-** Western Blot membranes of saliva samples collected from mosquitoes infected with ZIKV or CHIKV.

**Supplementary figure 7-** Comparative analysis of salivation performance and saliva infectivity from mosquitoes infected with ZIKV or CHIKV, in individual replicates.

**Supplementary figure 8-** Comparative analysis of salivation performance after mosquitoes feeding in mice.

**Supplementary figure 9-** Imaging of ZIKV and saliva reporters in whole mosquito mounts using the Fudgel transgenic line.

**Supplementary figure 10-** Analyzing the localization of ZIKV and saliva markers in salivary glands from Fudgel mosquitoes.

### Supplementary Tables

**Supplementary table 1-** Top 50 genes expressed in the distal lateral lobes (dll) of the salivary glands and associated dll averaged UMAP score.

**Supplementary table 2-** Salivary glands lobes-specifically expressed genes and associated abundance in saliva.

**Supplementary table 3-** Western Blot raw data for V5-mTurq signal detection by Western Blot.

**Supplementary table 4-** Western Blot raw data for V5-mTurq detection in saliva samples upon different physiological states.

**Supplementary table 5-** Salivary glands fluorescence quantification raw values and CTCF calculations for purging measurement after natural blood-feeding or forced salivation.

**Supplementary table 6-** Western Blot raw data for V5-mTurq detection in saliva samples upon ZIKV and CHIKV infection.

**Supplementary table 7-** qPCR raw data for ZIKV, CHIKV and mTurq RNA detection in mosquitoes carcasses.

**Supplementary table 8-** Primers used for Fudgel genetic construct cloning, transgene insertion mapping and qPCR.

**Supplementary table 9-** Mean area (µm2) comparison between mTurq clusters in Mock and ZIKV-infected individual salivary glands.

### Supplementary Video

**Supplementary video 1-** Live V5-mTurq detection in a Fudgel female proboscis upon forced salivation in PBS.

## Notes

### Competing Interest Statement

The authors have declared no competing interest.

### Summary of Updates

This version of the manuscript has been revised to confirm the results obtained in an artificial salivation system with in vivo data. We first confirmed that mosquitoes infected with ZIKV salivate significantly less while probing on mice, with similar feeding ratio. We also showed that ZIKV infection affects the organisation of saliva reporters in infected salivary glands, and that viral particles seem to displace saliva reporters in acinar cells, going along with the observed absence of correlation between virus and saliva release in forced salivation assays.

