## Supplementary figures for "Arboviruses disrupt salivary gland organization and decrease salivation in *Aedes aegypti* mosquitoes"

**Affiliations**
[1] Inserm U1257, CNRS UPR9022, University of Strasbourg, Strasbourg, France
[2] CNRS UPR3572, University of Strasbourg, Strasbourg, France
[3] Department of Biochemistry and Immunology, Instituto de Ciências Biológicas, Universidade Federal de Minas Gerais, Belo Horizonte/MG, Brazil
[4] CNRS UPR9022, University of Strasbourg, Strasbourg, France
[5] MIVEGEC, Université de Montpellier, IRD, CNRS, 34394 Montpellier, France

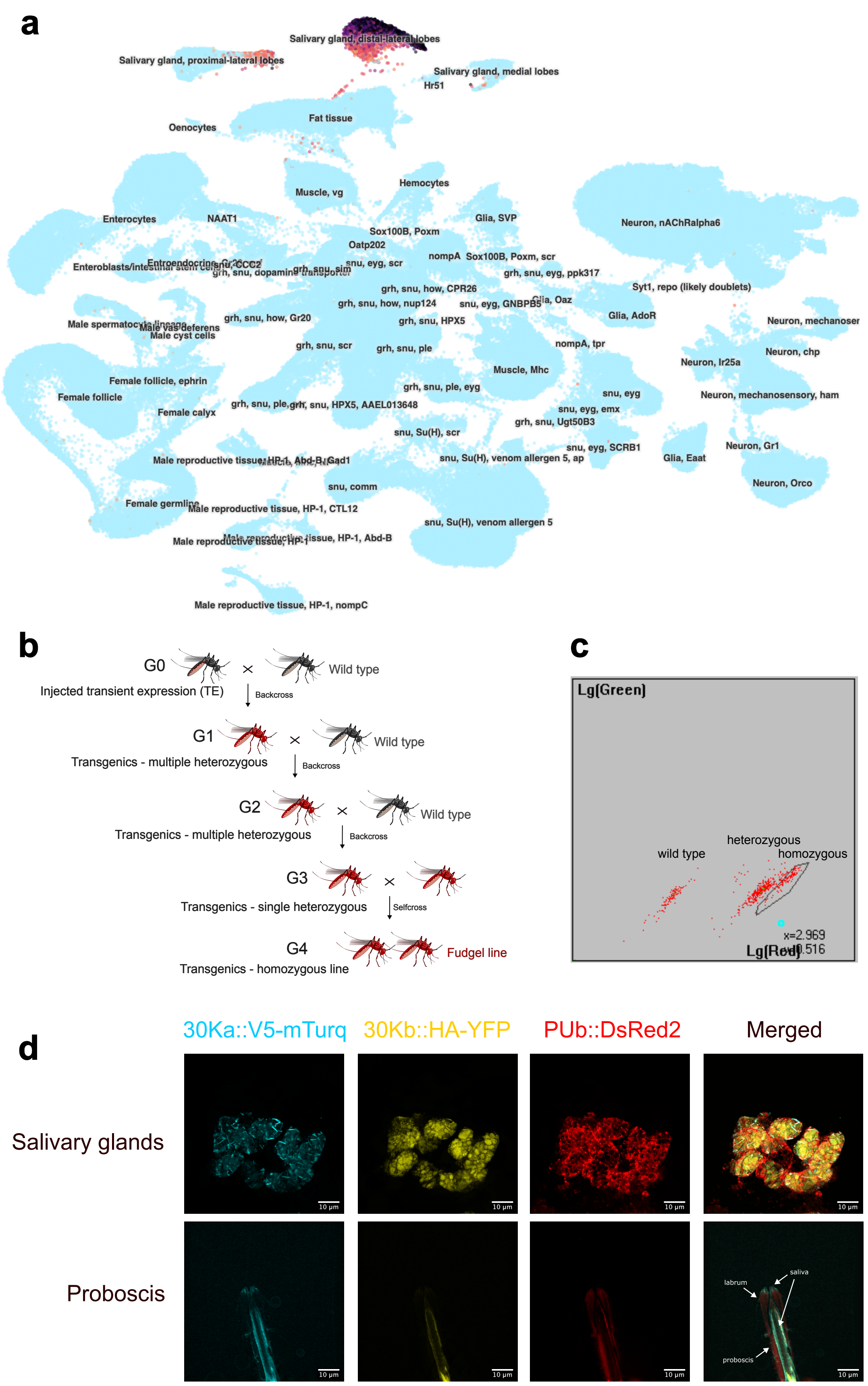


**Supplementary figure 1- Establishment and phenotypic characterization of the Fudgel transgenic line**
**a**) Combined UMAP representation of *30Ka* (*AAEL010228/SAAG-4*) and *30Kb* (*AAEL010235/aegyptin*) expression patterns in *Ae. aegypti* snRNAseq data from whole body generated with the UCSC Cell Browser (data from the Mosquito cell Atlas in Goldman et al. 2025). **b**) Crossing scheme for the establishment of the Fudgel line. **c**) Fluorescent profile of mixed G4 neonate larvae (shown in b) showing populations of wildtype, heterozygous and homozygous Fudgel mosquitoes. **d**) Confocal images of mTurq, YFP and DsRed2 in salivary glands and proboscis of Fudgel mosquitoes. Exposure time was 10-fold higher for YFP compared to mTurq. Z-stacks and merging were performed on ImageJ.

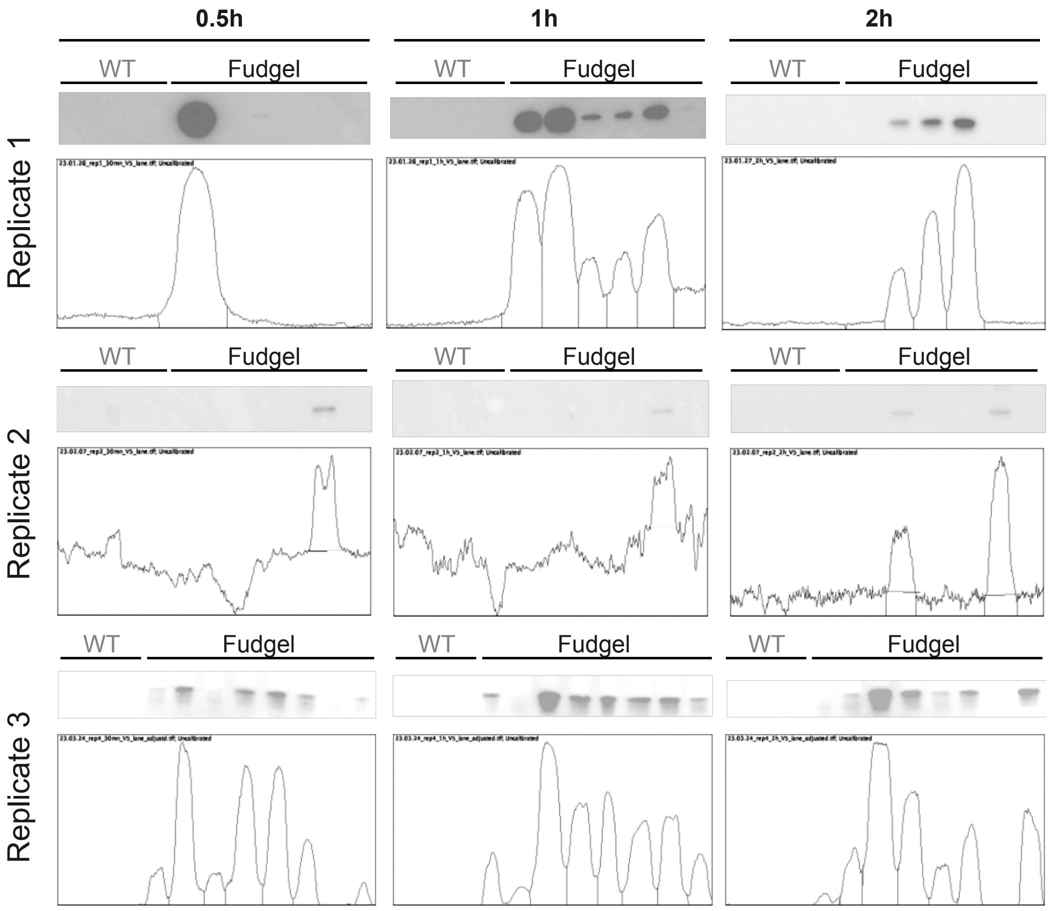


**Supplementary figure 2- Quantification of saliva reporter levels by Western Blot**

Western Blot anti-V5 and associated quantification plots for individual mosquito saliva samples collected after 0.5h, 1h or 2h of forced salivation for wild type mosquitoes (WT) or transgenic Fudgel mosquitoes (3 independent replicates). Quantification plot generated with ImageJ. Signal quantification data is shown in Supplementary table 3.

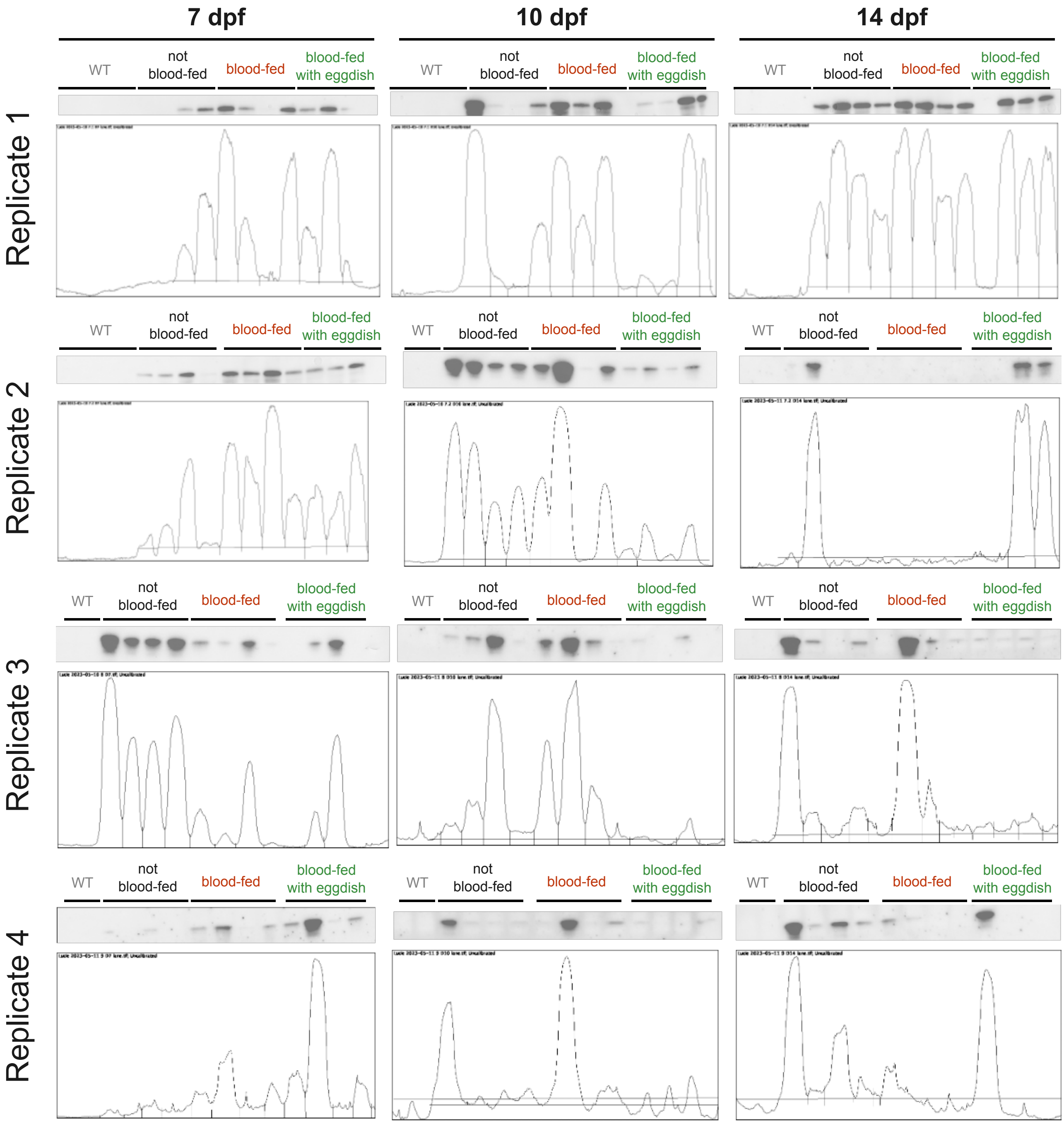


**Supplementary figure 3- Quantification of the saliva reporter in samples collected from mosquitoes in different physiological states**

Western Blot anti-V5 and associated quantification plots for individual mosquito saliva samples collected after 1h of forced salivation for wild type mosquitoes (WT) or transgenic Fudgel mosquitoes in different physiological states (not blood-fed, blood-fed or blood-fed with egg dish) (4 independent replicates). Quantification plot generated with ImageJ. Signal quantification data is shown in Supplementary table 4.

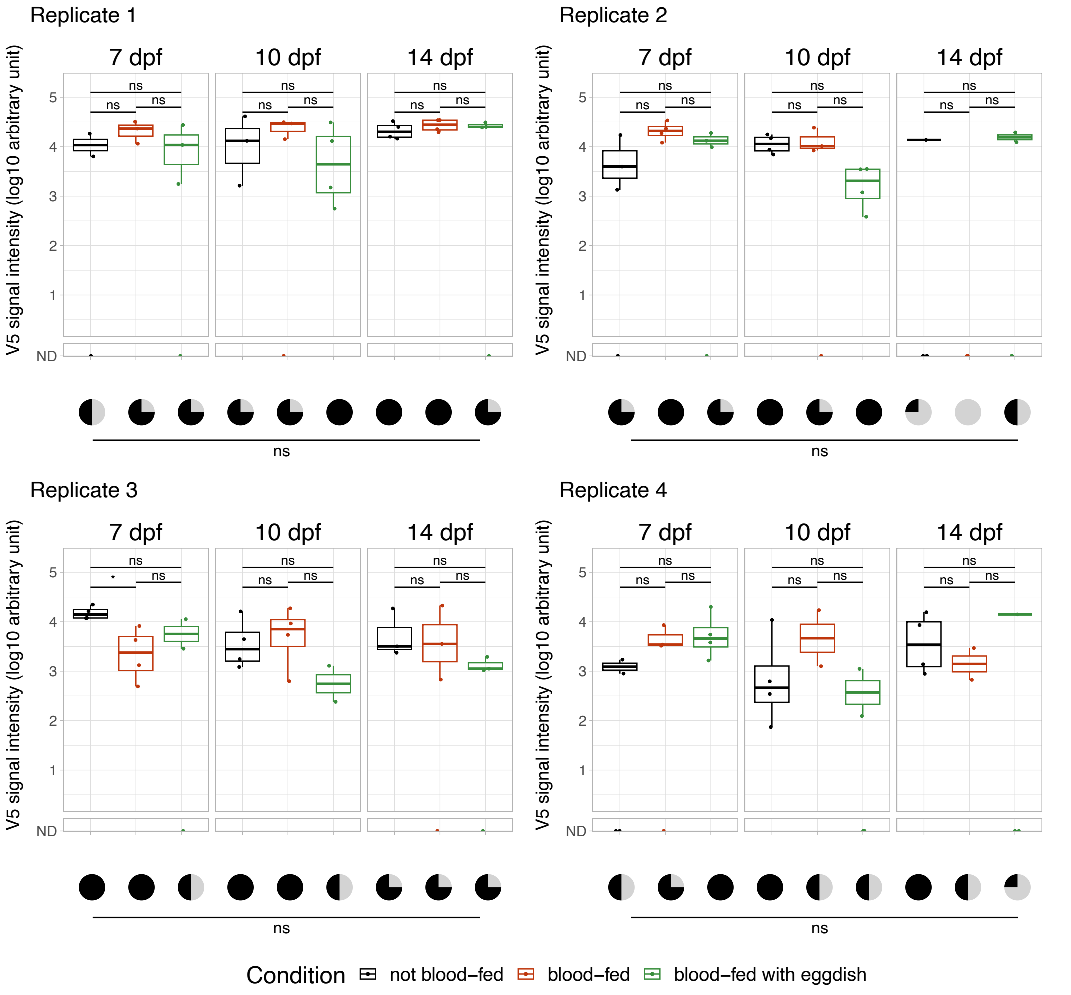


**Supplementary figure 4- Individual replicate plots for @V5-mTurq detection in saliva samples collected from mosquitoes in different physiological states**

V5 signal intensity in individual saliva samples of Fudgel mosquitoes in different physiological states after 1h of forced salivation (log10 scale). 4 independent replicates are shown. Comparison of signal intensity: Kruskall-Wallis test, threshold 0.05. Comparison of prevalence of salivation: Fisher's exact test, threshold 0.05.

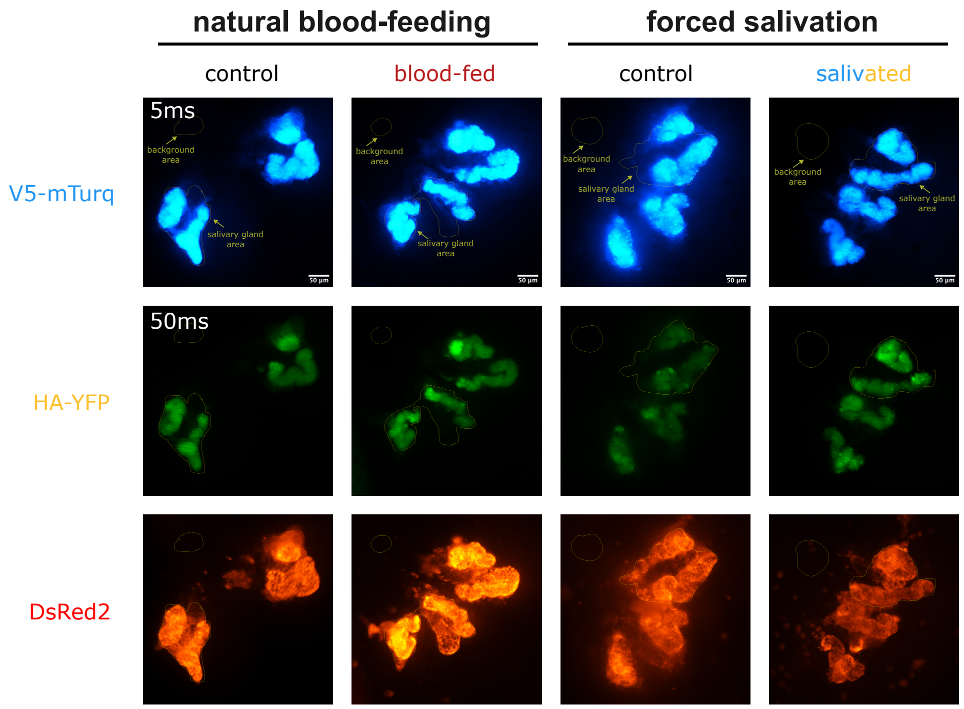


**Supplementary figure 5- Fluorescent images of salivary glands from Fudgel mosquitoes after natural blood-feeding or forced salivation**

Salivary glands (SG) fluorescence images for mTurq, YFP and DsRed2 from mosquitoes 1h after natural blood-feeding on mice or forced salivation. Exposure time was 10-fold higher for YFP compared to mTurq. Signal quantification data and CTCF calculations are shown in Supplementary table 5.

**a**

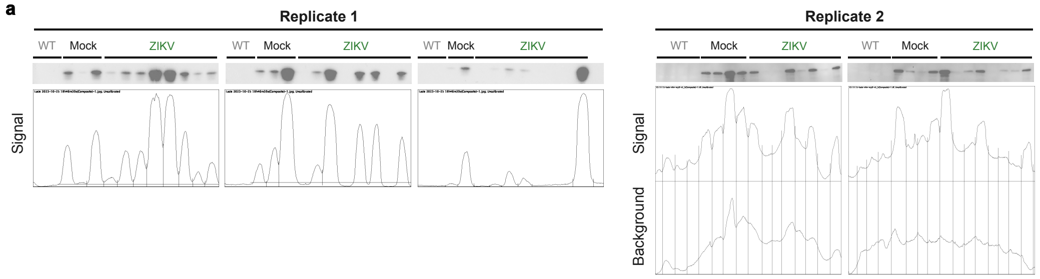


**b**

| Condition | Signal | Background | Difference |
|---|---|---|---|
| Mock | 23106.104 | 13587.983 | 9518.121 |
| Mock | 22327.497 | 15793.548 | 6533.949 |
| Mock | 44342.276 | 31376.518 | 12965.758 |
| Mock | 31580.619 | 26390.74 | 5189.879 |
| Mock | 34507.175 | 22013.347 | 12493.828 |
| Mock | 17421.619 | 16527.255 | 894.364 |
| Mock | 17480.841 | 14878.134 | 2602.707 |
| Mock | 28820.154 | 17148.861 | 11671.293 |
| ZIKV | 28057.518 | 25223.225 | 2834.293 |
| ZIKV | 11343.841 | 13543.305 | -2199.464 |
| ZIKV | 12634.719 | 12937.648 | -302.929 |
| ZIKV | 35757.69 | 25638.104 | 10119.586 |
| ZIKV | 16454.426 | 17569.548 | -1115.122 |
| ZIKV | 21836.033 | 17337.205 | 4498.828 |
| ZIKV | 3227.619 | 9129.619 | -5902 |
| ZIKV | 13641.326 | 10003.669 | 3637.657 |
| ZIKV | 42720.397 | 18570.033 | 24150.364 |
| ZIKV | 15050.497 | 13616.669 | 1433.828 |
| ZIKV | 16046.376 | 14855.083 | 1191.293 |
| ZIKV | 28024.569 | 18187.276 | 9837.293 |
| ZIKV | 10141.426 | 11902.719 | -1761.293 |
| ZIKV | 10148.012 | 10225.891 | -77.879 |
| ZIKV | 8422.569 | 10944.326 | -2521.757 |
| ZIKV | 14217.447 | 4867.326 | 9350.121 |

**c**

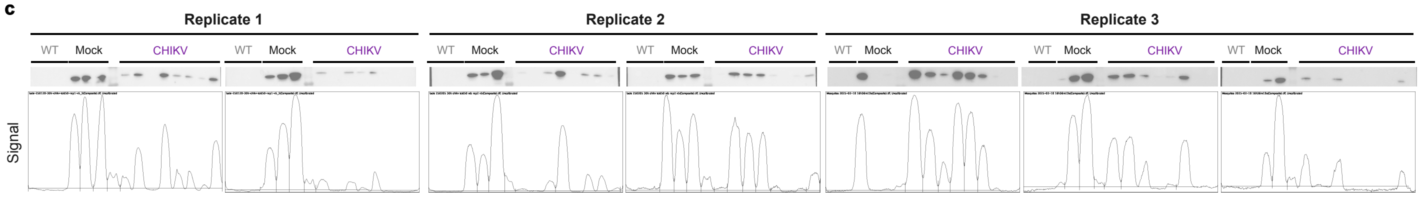


**Supplementary figure 6- Western Blot membranes of saliva samples collected from mosquitoes infected with ZIKV or CHIKV**
Western Blot @V5-mTurq and associated quantification plots for individual mosquito saliva samples infected with ZIKV 14 dpi (**a-b**) or CHIKV 9 dpi (**c**), for each experimental replicate. For the second replicate of ZIKV, background removal had to be performed to quantify western blot bands intensity (**b**). Quantification plot generated with ImageJ. Signal quantification data displayed in Supplementary table 6.

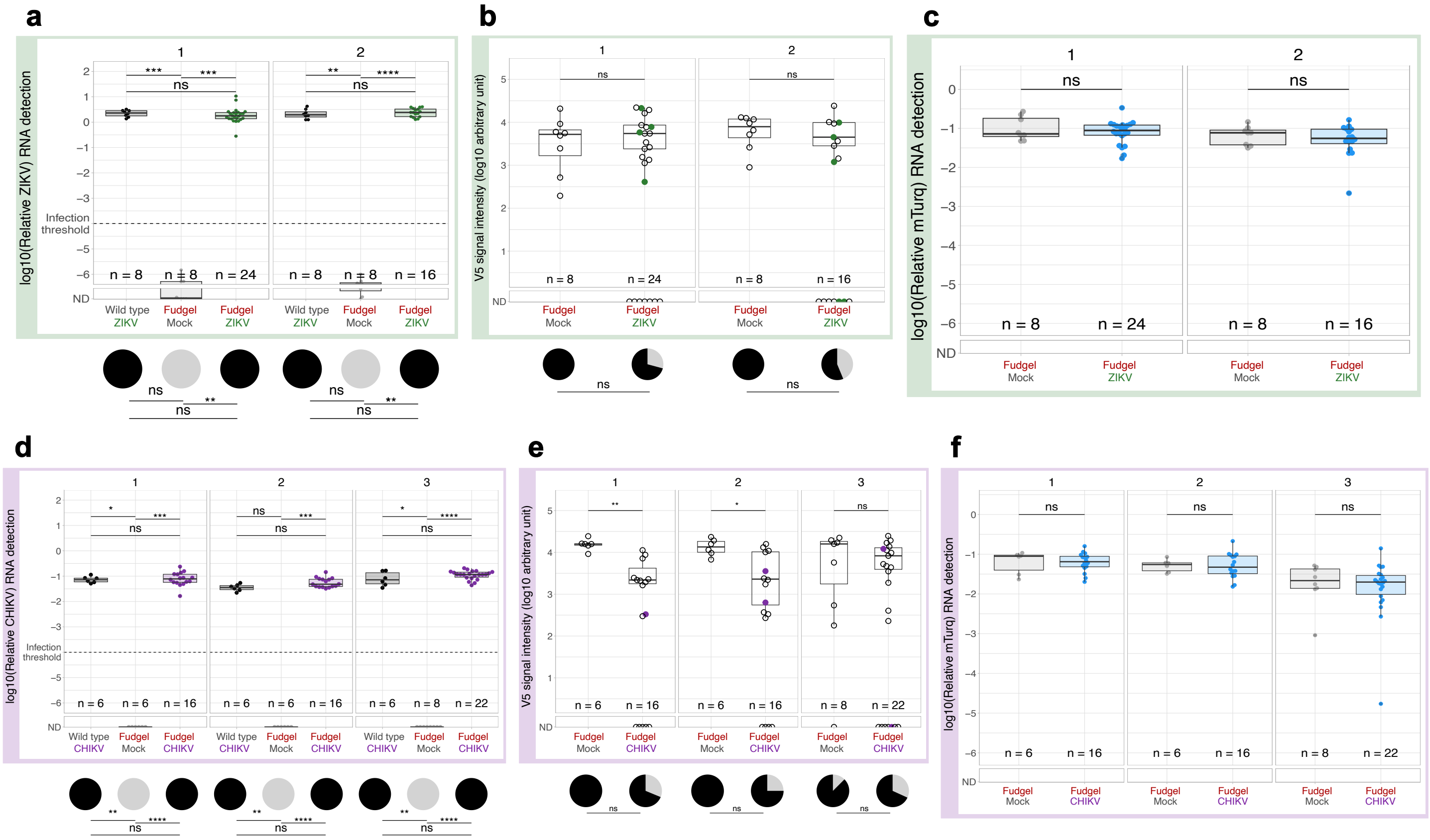


**Supplementary figure 7- Comparative analysis of salivation performance and saliva infectivity from mosquitoes infected with ZIKV or CHIKV, in individual replicates**

**a**) Detection of ZIKV by RT-qPCR in mosquitoes at 14 days post infection for each experimental replicate. **b**) V5 signal intensity in individual samples of control or ZIKV-infected mosquitoes for each experimental replicate. Saliva samples associated with the presence of infectious virus are marked in green. **c**) Detection of V5-mTurq expression by RT-qPCR in control and ZIKV-infected mosquitoes for each experimental replicate. **d**) Detection of CHIKV by RT-qPCR in mosquitoes at 9 days post infection for each experimental replicate. **e**) V5 signal intensity in individual samples of control or CHIKV-infected mosquitoes for each experimental replicate. Saliva samples associated with the presence of infectious virus are marked in purple. **f**) Detection of V5-mTurq expression by RT-qPCR in control and CHIKV-infected mosquitoes for each experimental replicate. Comparison of viral loads in carcasses, V5 signal intensity and V5-mTurq levels in carcasses: Kruskall-Wallis test, threshold 0.05. Comparison of prevalences: Fisher's exact test, threshold 0.05.

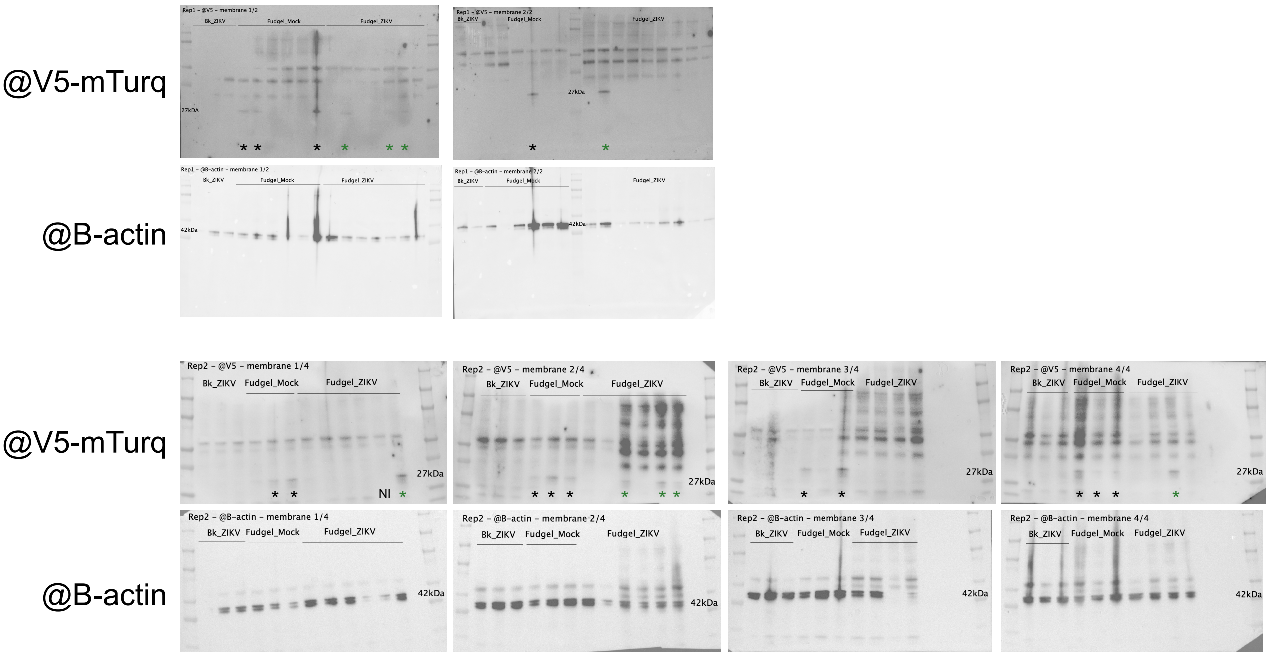


**Supplementary figure 8- Comparative analysis of salivation performance after mosquitoes feeding in mice**
Western Blot @V5-mTurq (salivation reporter) and @B-actin (loading control) for mice biopsy samples bitten by individual mosquitoes, wild type (WT) or transgenic Fudgel, infected (ZIKV) or non-infected with ZIKV (Mock) in 2 independent replicates. *:positive for V5-mTurquoise detection.

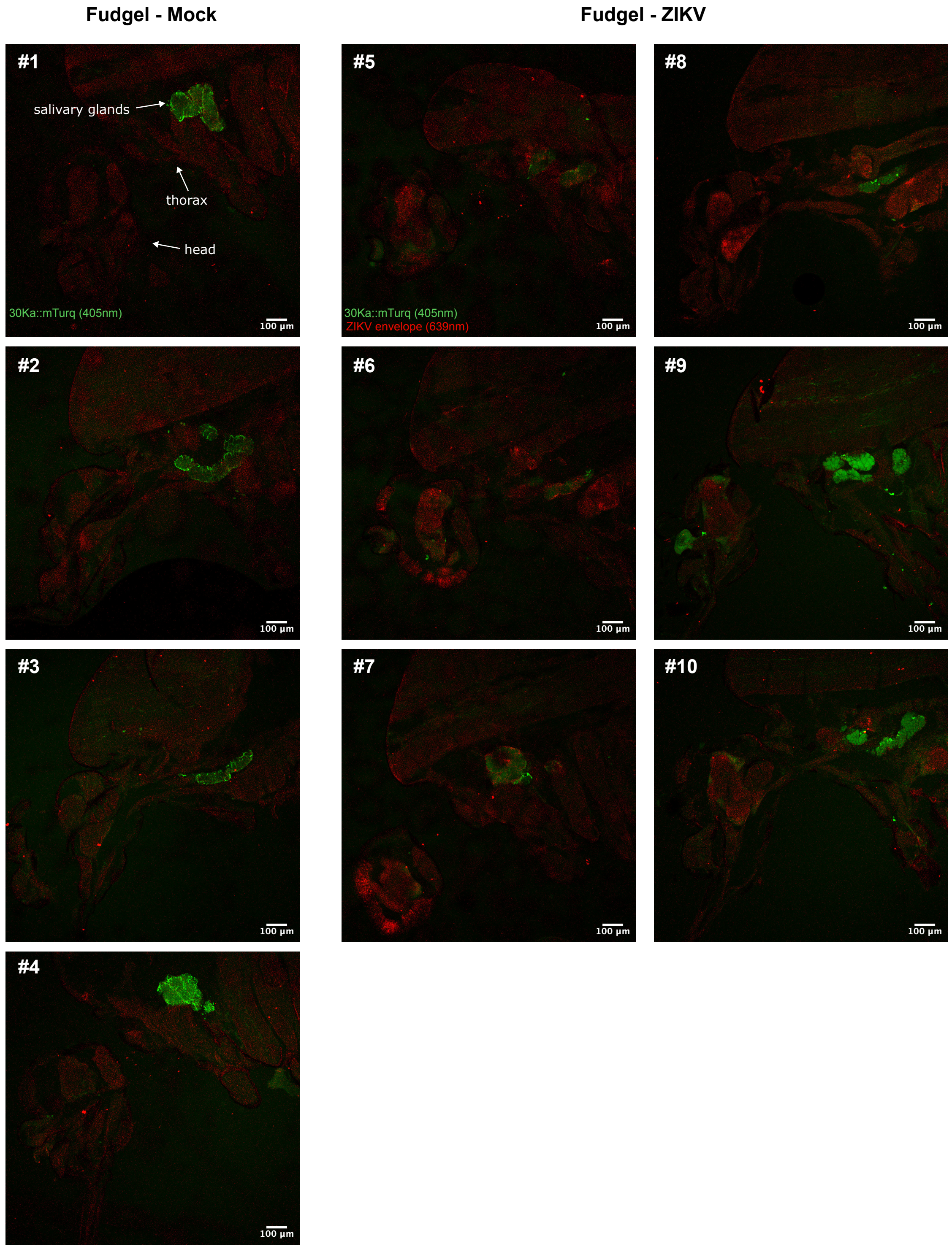


**Supplementary figure 9- Imaging of ZIKV and saliva reporters in whole mosquito mounts using the Fudgel transgenic line**
10x confocal images of whole Fudgel mosquitoes, either non-infected (Mock) or infected with ZIKV (ZIKV). Whole mosquitoes 14 dpi were embedded in OCT and microtome-sliced (30µm thickness). ZIKV virus was stained using a 4G2 antibody, secondary Alexa 647, and images were acquired using a confocal Zeiss, Beam Path LSM980 microscope.

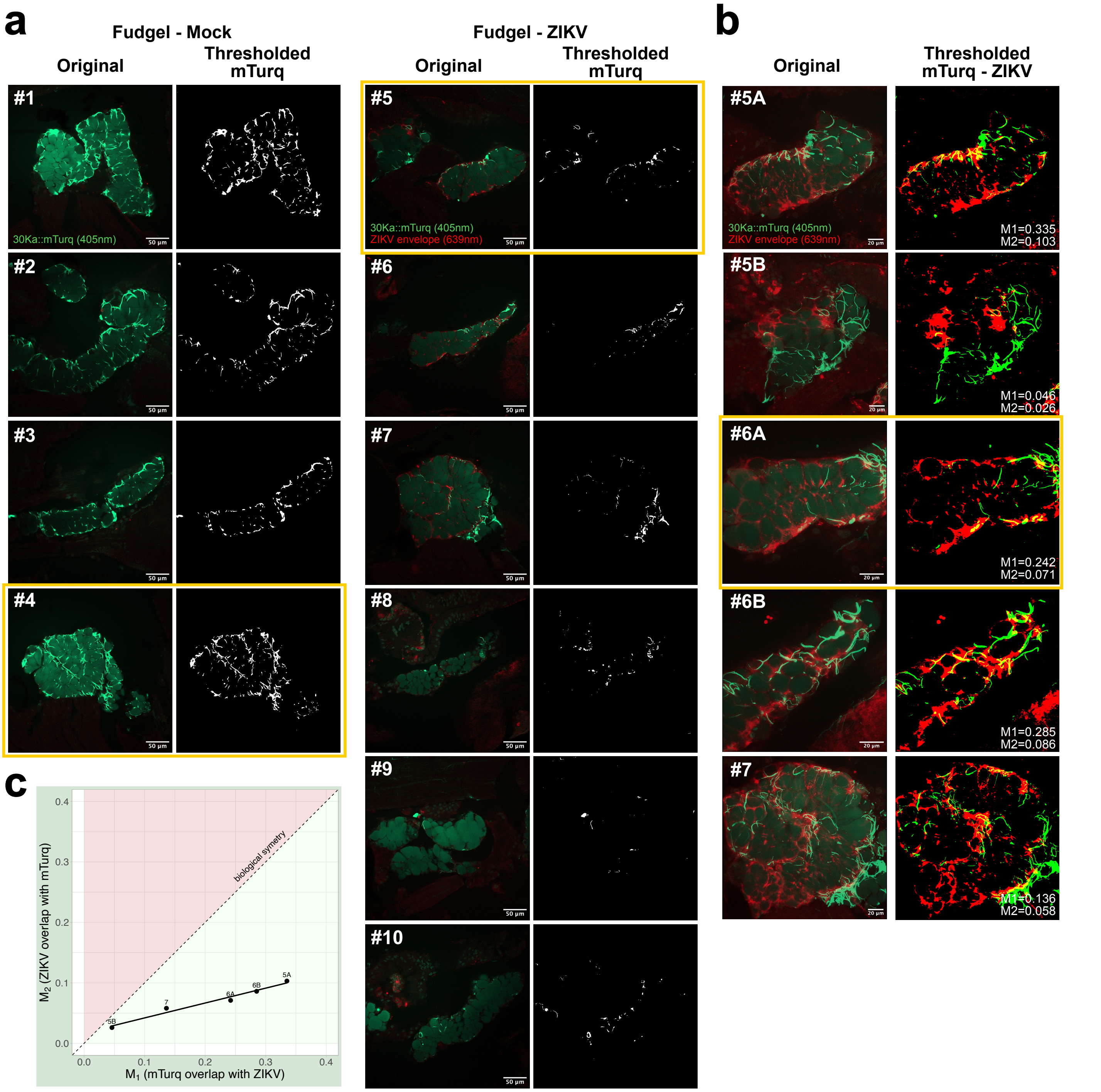


**Supplementary figure 10- Analyzing the localization of ZIKV and saliva markers in salivary glands from Fudgel mosquitoes.** **a**) Confocal images (40x) of Mock (#1-4) and ZIKV-infected salivary glands (#5-10): mTurq (green) and the ZIKV E protein are shown (red). Confocal images ("Original") were deconvolved using the "Iterative Deconvolve 3D" plugin in ImageJ (20 iterations)[40] and saliva aggregates were highlighted using the "2D Threshold" tool on the "Mitochondria Analyzer" plugin ("Thresholded")[41]. **b**) 3D projections (63x) of ZIKV-infected salivary glands (#5-7), obtained with the "3D Project" tool in ImageJ. Stacks were deconvolved using the "Iterative Deconvolve 3D" plugin in ImageJ (20 iterations), thresholded using a Otsu method in ImageJ and Manders' overlap coefficients were obtained using the "JaCoP" plugin in ImageJ[43]. Images displayed in Figure 5 are highlighted in yellow.
