## Supplementary figures and images for "Arboviruses disrupt salivary gland organization and decrease salivation in *Aedes aegypti* mosquitoes"

### Supplementary video

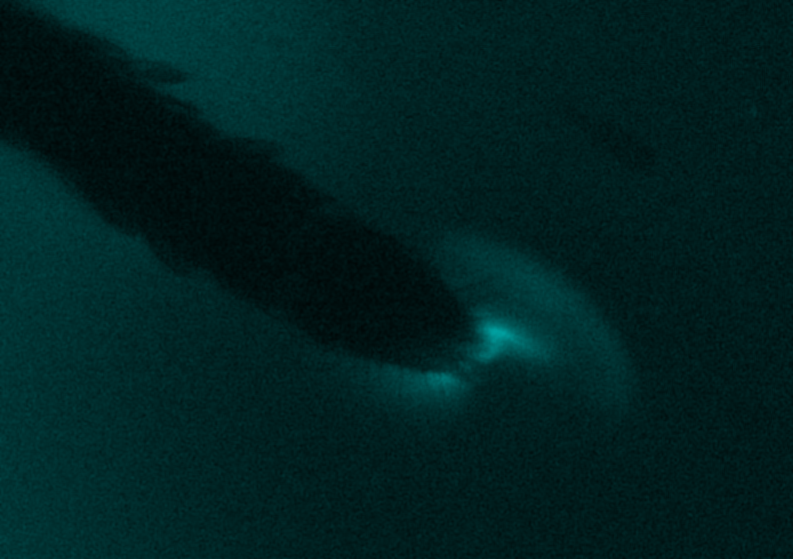
